# LAMP sheds light on how sequence to function models learn variant effects during post-training

**DOI:** 10.64898/2026.11.13.766567

**Authors:** Shiron Drusinsky, Katherine S. Pollard

## Abstract

Genomic sequence-to-function (S2F) models show promise for interpreting noncoding genetic variation, yet they currently underestimate the effects of most causal variants. This leads to fine-mapping failures that are reflected in their underperformance relative to linear baselines at predicting gene expression across individuals from personal genome sequences. Improving personal genome predictions, which reflect this underestimation, is therefore critical. There is broad interest in post-training S2F models to improve these predictions, yet many groups have shown that doing so with personal genome sequences and paired expression values does not meaningfully improve predictions on held-out genes and people (HOGP). Why these post-training efforts to improve predictions on HOGP, and therefore causal variant underestimation, remains unclear. To understand why, we introduce LAMP (Learning of Artificial Motif Perturbations), a framework in which artificial SNVs (aSNVs) perturbing transcription factor motifs are substituted into personal genome datasets genome-wide with large, consistent, and causal effects on gene expression. These aSNVs are designed to be deliberately easy to learn, yet fine-tuning Enformer using this LAMP framework led to persistent underperformance on HOGP, which suggests fundamental challenges for learning variant effects during post-training that are independent of the complexity of natural genetic variation. Our findings suggest this underperformance is caused in part due to challenges learning to strengthen enhancer-gene links during post-training. In particular, models struggle with learning to rewire attention weights to relate aSNV detection, over distance, to a change in expression of the target gene. This limitation is caused by strong pre-trained priors that downweight distal regulatory elements, causing extreme data inefficiency in learning to attend to distal elements, and variants within them, during post-training. Second, sequence variability between loci containing different instances of the same TF motif obscures otherwise easy-to-learn motif perturbation patterns and also hinder learning of variant effects. Our results suggest that the scale of observed human genetic variation is vastly insufficient to maximize S2F model accuracy on HOGP via simple fine-tuning, consistent with past outcomes. We propose data augmentation strategies that, under LAMP, significantly improve learning of aSNV effects by strategically targeting the limitations we identified. We release LAMP as a tool to troubleshoot and optimize future S2F post-training objectives.

## Introduction

Genomic sequence-to-function (S2F) models take only DNA sequence as input, and accurately predict context-specific outcomes for assays that measure mechanisms of gene regulation, including DNA accessibility, transcription factor binding, histone post-translational modifications, and gene expression^1–8^. A subset of S2F models, including Enformer^1^, Borzoi^2^, and AlphaGenome^3^, can integrate information from distal regulatory elements to form gene expression predictions, including predictions of how variants within those elements alter expression. This class of long-range S2F models (hereafter, S2F models) is the focus of this study. Since these S2F models predict regulatory activity for any input sequence, they can be applied to haplotypes with complex variant interactions and to rare variants. They therefore represent tremendous potential for fine-mapping and interpreting noncoding variation, which is critical for decoding the gene regulatory code and learning disease mechanisms^7,9^.

To date, however, S2F models have shown moderate to poor performance on benchmarks that evaluate genetic variation not seen during model training. First, S2F models pervasively underestimate the gene expression effects of causal single-nucleotide variants (SNVs) relative to nearby non-causal ones^10^. Second, this underestimation causes inaccurate gene expression predictions from personal genome sequences^10–13^, which aggregate predicted effects across every variant an individual carries at a locus and therefore depend on causal variants being weighted above their neighbors. This makes personal expression prediction an important implicit test of fine-mapping. Third, underestimation is most common among variants distal to the transcription start site (TSS)^10^, owing to S2F models’ neglect of the effects of distal regulatory elements on gene expression^1–3,14^, and this distal insensitivity leads to personal genome prediction errors.

Each successive generation of S2F models has brought only small improvements on the critical personal genome and underestimation issues^11–13^, suggesting the standard reference genome pre-training paradigm is unlikely, on its own, to resolve them. Because this pre-training excludes genetic variation, there has been great interest in incorporating variation during post-training, whether by training on variant effects^15^ or on paired personal genome and gene expression datasets^16–19^. These attempts have been most successful at improving variant effect predictions for held-out promoter-proximal variants^15^, but have led to weak or no improvements on personal expression predictions from larger genomic windows surrounding held-out genes that contain held-out variants^16–19^. Why post-training helps the former but not the latter remains unclear. Because underestimation of distal variant effects is a major contributor to personal genome prediction errors^10^, one possibility is that post-training on genetic variation datasets is more effective for improving local rather than distal variant effect predictions, though this has not been clearly demonstrated.

There remains significant room for improving S2F predictions for causal variants and personal genomes. To do so, it is critical to first clearly understand the sources of expression prediction failures. In this study, we introduce a framework to solve this problem, named LAMP (Learning of Artificial Motif Perturbations). Specifically, we design clear and causal, yet completely artificial, gene expression-modifying SNVs that act by perturbing the same transcription factor’s (TF) motif throughout the genome, yielding a stereotyped and therefore easily learnable pattern, and introduce them into paired personal genome and gene expression datasets to study the ability of S2F models to learn their effects during fine-tuning. The LAMP approach enabled us to train on far more causal artificial SNVs (aSNVs) than the number of causal SNVs natural datasets are expected to contain, to know their identities with certainty, and to systematically vary their properties in a controlled manner to carefully probe how S2F models learn their causal effects.

By simplifying the learning task, our study identifies fundamental failure modes that hinder S2F learning of variant effects and that would otherwise be extremely difficult to disentangle from the complexity of real genetic variation. One obstacle is data inefficient rewiring of attention weights to link distal regulatory elements to target genes – a consequence of strong, miscalibrated pre-trained priors that bias S2F models away from distal regulatory elements – which hinders learning and generalizing variant effects beyond the promoter. A second is sequence variability between loci containing different instances of the same TF motif, an underappreciated challenge inherent to the genome that obscures otherwise easy-to-learn motif perturbation patterns. Together, these issues hinder post-training of variant effects to such an extent that human genetic variation datasets are unlikely to be sufficient to optimize S2F models on fine-mapping and personal genome prediction tasks via simple fine-tuning. Instead, we propose data augmentation strategies that significantly improve learning of aSNV effects by strategically targeting these limitations. We also speculate that strategically designed perturbational datasets (e.g., CRISPRi-FlowFISH^20^) that disproportionately sample distal elements and sample each motif deeply enough to overcome sequence context variability will help bridge this gap. We propose LAMP as a tool to troubleshoot and optimize these post-training strategies.

## Results

### LAMP (Learning of Artificial Motif Perturbations) generates artificial, causal expression modifying variants to study how S2F models learn their effects

LAMP is a framework for generating clear and causal, yet completely artificial, gene expression-modifying SNVs and introducing them into S2F training datasets as a means to study the ability of S2F models to learn variant effects on gene expression. These artificial SNVs (aSNVs) are designed to have two key properties: (1) They confer a strong, independent change to gene expression; and (2) they act via a consistent, unambiguous mechanism – perturbing the same transcription factor’s (TF) motif in multiple loci genome-wide, yielding a stereotyped association. We therefore expected aSNVs to be much easier for a model to learn relative to natural variants, while still mirroring a TF motif-perturbing mechanism by which natural noncoding regulatory variants may act.

To effectively track how S2F models learn variant effects, LAMP enables the controlled modification of causal variant parameters that are unknown in observational data, such as: effect size, minor allele frequency (MAF), distance to the TSS, number of causal aSNVs per locus or total in the training dataset, and choice of aSNV-perturbed TF motif(s). Simulated aSNVs can be injected into the training data, validation data, and/or test data, and the variant parameters need not be the same in each set. Hence, LAMP is a highly flexible approach to test the ability of any S2F expression predicting model to detect different types of causal variants.

In this study, we mostly used LAMP to modify personal genome datasets, consisting of paired whole genome sequences (WGS) and RNA-seq, to diagnose why past efforts to post-train S2F models with these datasets only minimally improved unseen variant effect predictions around held-out genes^16–19^. As LAMP requires only DNA sequence inputs (to introduce aSNVs) and gene expression outputs (for concomitant expression changes), it can also be applied to other dataset types. Here, we demonstrate this by using LAMP to generate *in silico* mutagenesis (ISM) datasets where examples consist of reference genome sequence with either of the two alleles substituted and then test whether this input data format, which isolates the effect of a single variant, changes how readily expression differences are learned during fine-tuning.

### Using LAMP to introduce controlled artificial SNVs into personal genome datasets reveals inefficient learning of variant effects

To better understand why past attempts to improve S2F personalized gene expression predictions have largely failed^16–19^, we attempted to fine-tune Enformer with a dataset of paired WGS and “Whole Blood” gene expression values (n=670 individuals) from the Adult Genotype-Tissue Expression (GTEx) Project^21^ (**Methods**). To simplify the learning task and isolate specific failure modes, we used LAMP to introduce G/C substitutions at the 6th position of one naturally occurring STAT1 motif instance (JASPAR^22^ motif MA0137.4) per gene, assigning either the G or C allele to different personal genomes, along with strong, consistent gene expression increases associated with the G allele (the “effect allele”; **Fig. 1, Methods**). Other motifs are explored after deeply investigating STAT1.

**Figure 1:**
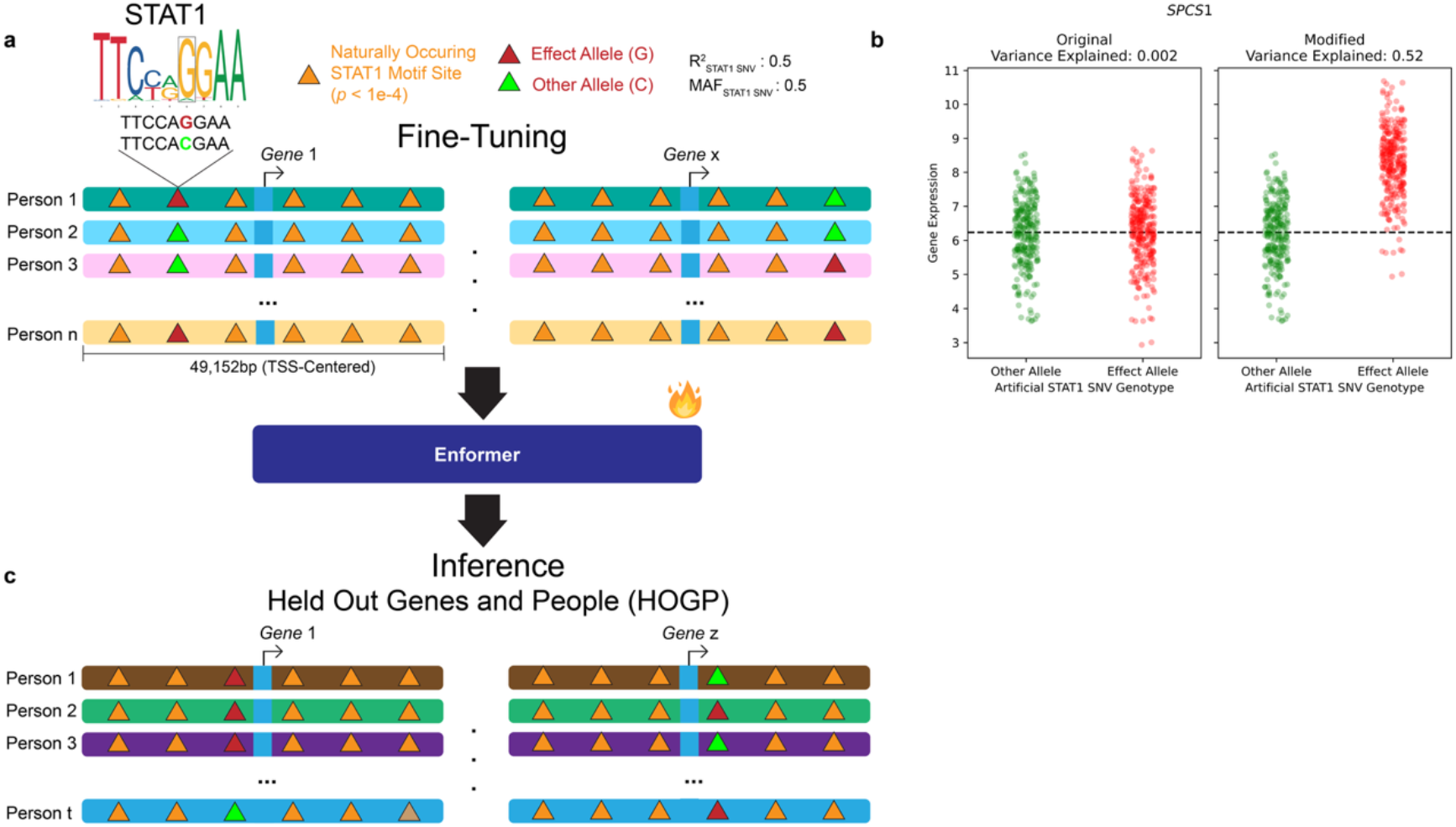
Schematic describing the LAMP framework. **a**) The aSNV perturbs one naturally occurring motif (orange triangle) around each gene. In a given experiment, the aSNV perturbs the same TF’s motif genome-wide (here, STAT1). Each individual is randomly assigned either the effect allele (red, G) or the other allele (green, C). **b**) Individuals assigned the effect allele have their original expression value for a given gene increased, such that the aSNV genotype explains 50% of the variance in gene expression across individuals for the gene (the change for *SPCS1* is shown as a random example; **Methods**). Together, this amounts to a strong signal (large effect to expression) and a clear learnable pattern (the aSNV perturbs the same motif genome-wide, with the same two alleles, in the same relative position, directionally consistent effects, and random allele assignment ensures it is not in LD with natural SNVs to make its causal effect unambiguous) for Enformer to learn during fine-tuning. **c**) We then evaluated Enformer on held-out genes and people (HOGP), perturbed by the aSNV in the same way, and evaluated the extent to which Enformer can learn this idealized variant effect pattern and generalize it to new contexts. This reflects how we expect S2F models to learn motif perturbation patterns, caused by variants, during post-training, but the setup is simplified and idealized to ensure shortcomings cannot be attributed to the complexity of natural genetic variation.

Unless otherwise stated, the STAT1 aSNV effect allele is assigned to each personal genome at random at a 50% MAF and is designed to explain 50% of the variance in each gene’s expression across individuals (**Fig. 1**, **Methods**). This amount of gene expression heritability is much greater than what is found in most loci, even when considering the combined effects of natural variants (**Supp. Fig. 1a**). The aSNV therefore has an effect size outsized relative to natural variants, and is the predominant genetic driver of expression at any locus where it is placed (**Supp. Fig. 1b**). We engineered the aSNV this way to incentivize Enformer to learn its effects during fine-tuning. To support a consistent pattern of aSNV-caused expression changes to different genes, we standardized cross-individual variance in gene expression so that the aSNV explains the same fraction (50%) of expression variance with the same effect size for every gene (**Methods**). Together, this models the scenario of one causal, very common, and very strong eQTL-like SNV acting via the STAT1 motif perturbation mechanism around each gene, with the same outsized expression effect each time. Random allele assignment is expected to leave no other SNV in linkage disequilibrium (LD) with the aSNV, making its causal status unambiguous. The entire perturbation is a single nucleotide per gene, along with a concomitant gene expression change, with the background genomic sequences and baseline expression values controlled. Altogether, the large effect size, high MAF, stereotyped motif disruption, and lack of LD constitute an extremely detectable association relative to real causal variants; failure would therefore suggest fundamental limitations on learning variant effects rather than difficulty learning from the complexity of natural genetic variation.

We chose STAT1 for our initial experiments because (1) Most positions in the motif’s position weight matrix (PWM) are dominated by a preference to one nucleotide (**Fig. 1**). This ensures that genomic windows with significant matches to the PWM will be easily recognizable and therefore should be easier to learn by the model as a recurrent pattern. (2) Sequences significantly matching the STAT1 consensus PWM are prevalent and uniformly distributed within 49kb TSS-centered windows (**Supp. Fig. 2a-b**; Median of 16 STAT1 sites per gene). Rather than needing to alter DNA sequences to create a new motif, we can therefore choose among many naturally occurring motifs to place the aSNV into, based on which best addresses a particular experimental question. (3) STAT1 is expressed in GTEx “Whole Blood”, matching our gene expression dataset. Finally, (4) the STAT1 motif is palindromic, which we expected would support the model’s ability to learn the causal pattern, as the motif will appear nearly identical when either sequence orientation is passed into the model during training, thereby effectively doubling its sample size when training with random reverse complementation, a standard augmentation strategy. In our experiments, we applied LAMP perturbations to only highly significant STAT1 motif matches (*p* < 10^-4^; **Methods**), which we found to be a stringent cutoff that selected only 4.7% of the most easily recognizable motif instances (**Supp. Fig. 2c**). We note that these matching motifs may vary slightly in sequence from one another, yet all significantly match the consensus PWM. We chose not to restrict LAMP to exact STAT1 consensus sequence matches, so our experiments reflect how natural causal variants may perturb non-consensus motif-matching sequences.

**Figure 2:**
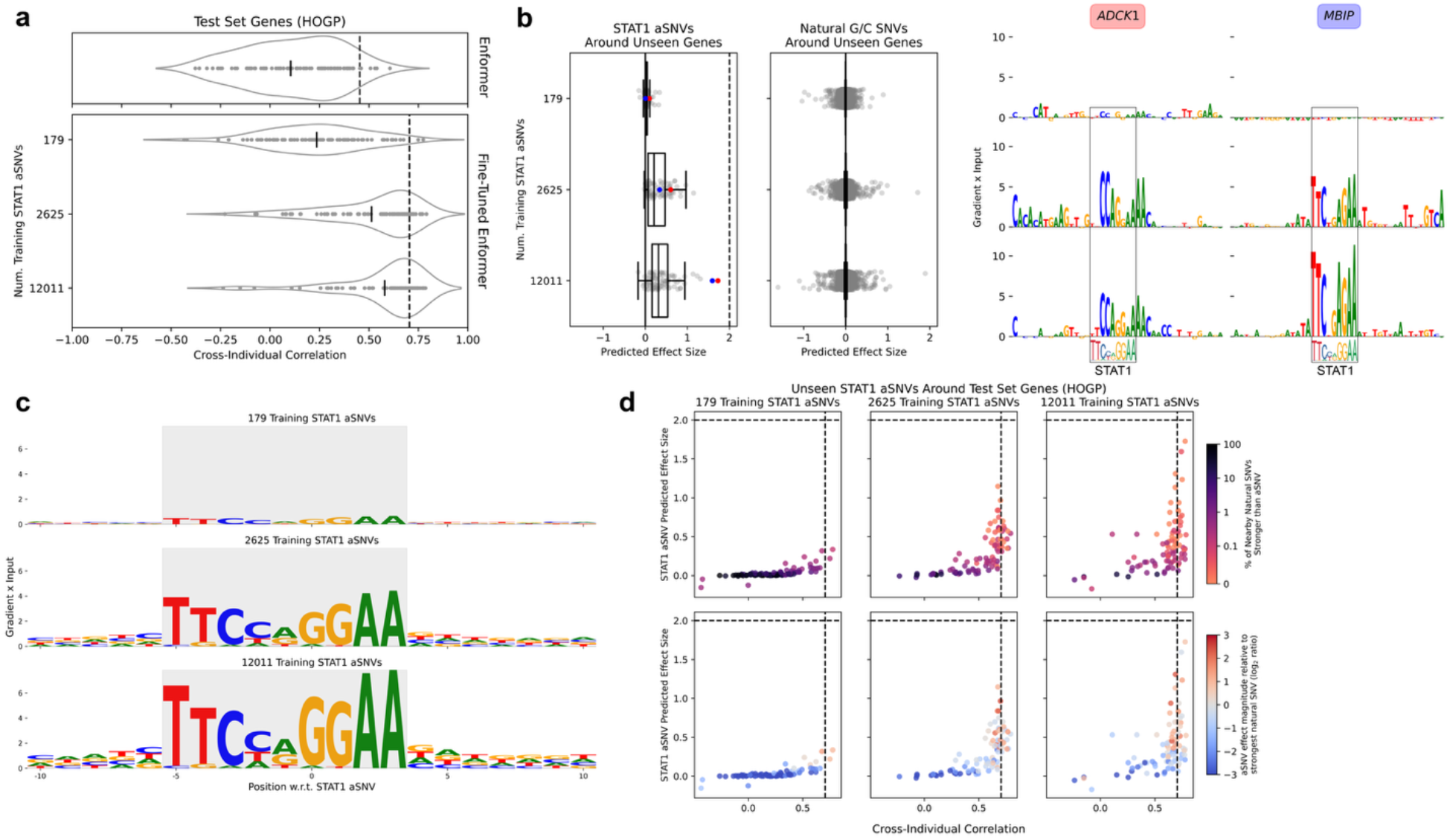
Enformer inefficiently learns idealized aSNV effects. **a**) Performance of fine-tune Enformer models on 88 test set genes and held-out people (HOGP), as measured by cross-individual correlation (bottom). Rows correspond to training dataset sizes. Dashed line corresponds to a linear baseline fit and evaluated on the aSNV genotype using the test set genes directly (**Methods**), marking the expected performance of a model that learns the gene expression differences the aSNV confers. On top are results from Enformer before fine-tuning, evaluated on the same HOGP, but that do not include the aSNV. The elastic net baseline (dashed line) is fit and evaluated on SNV genotypes that do not include the aSNV. Below are results after fine-tuning on the aSNV dataset, with the elastic net baseline fit using aSNV genotypes (**Methods**). aSNVs are placed into one random STAT1 motif around each gene. Relative to before pre-training, fine-tuning on personal genome datasets that include the aSNV leads to its improved generalization to HOGP, but even a model trained on the largest possible dataset underperforms the linear baseline. Each point is a different gene, with the cross-individual correlation averaged over three model replicates (different seeds), except for the 12,011 aSNV model, for which only two model replicates were trained to conserve resources. **b**) Left: predicted effect sizes for aSNVs from the same HOGP as in (**a**), with the true effect size marked by the dashed line; aSNV predicted effects are consistently underestimated, but increase with training dataset size. Middle: predicted effect sizes for all of the natural SNVs with G/C alleles around the same genes within the same 49kb window. Predicted effect sizes are averaged over model replicates as in (**a**). Right: Gradient x Input attributions (**Methods**) for STAT1 motifs perturbed by aSNVs marked by color in the leftmost panel. Increased aSNV predictions are not caused by increased predicted effect for arbitrary SNVs with the same G/C alleles. Instead, Enformer learns to detect the aSNV pattern by noticing its placement within the STAT1 motif. **c**) Gradient x Input attributions averaged over all candidate STAT1 motif instances around genes in the test set, including those that did not receive the aSNV, plus surrounding sequence. **d**) Predicted effect size for STAT1 aSNVs around test set genes (Y-axis) versus cross-individual correlations achieved for the same genes (HOGP), for models trained on different amounts of training data (columns). Points are colored by the percent of naturally occurring SNVs in each gene’s sequence window with a stronger predicted effect than the aSNV (top) or the log2 ratio of the aSNV’s predicted effect magnitude to that of the strongest natural SNV in the same window (bottom; values are clipped between −3 and 3 for axis visibility). Genes with more accurate HOGP predictions have aSNVs in the region with stronger predicted effects and that stand out more prominently above naturally occurring SNVs. Consistent with past findings^10^, this suggests weak HOGP predictions are a symptom of poor aSNV fine-mapping.

We tested the extent to which Enformer could learn how the STAT1 aSNVs perturb the STAT1 motif to increase expression from the instances it encounters around train genes, and then generalize that pattern to held-out genes and people (HOGP). This mirrors how we would expect Enformer to learn the more subtle effects of natural genetic variation from unaltered personal genomes. We first introduced the STAT1 aSNV into a single, randomly chosen pre-existing STAT1 motif around each train and held-out gene, and confirmed that the engineered expression change induced by the aSNV explains 50% of expression variance for each gene (**Supp. Fig. 3**). After fine-tuning Enformer on the STAT1 LAMP dataset, we evaluated it on HOGP. Relative to the scenario where pre-trained Enformer is evaluated on the same HOGP without the aSNV, HOGP accuracy increased. This suggests Enformer learned to generalize detection of the STAT1 aSNV to held-out parts of the genome after training on personal genome sequences that included similar STAT1 aSNV perturbations (**Fig. 2a**). Indeed, we found that the fine-tuned Enformer learned to detect this SNV by correctly noticing its placement within the STAT1 motif, rather than by associating arbitrary G/C substitutions with expression increases (**Figs. 2b-c**).

**Figure 3:**
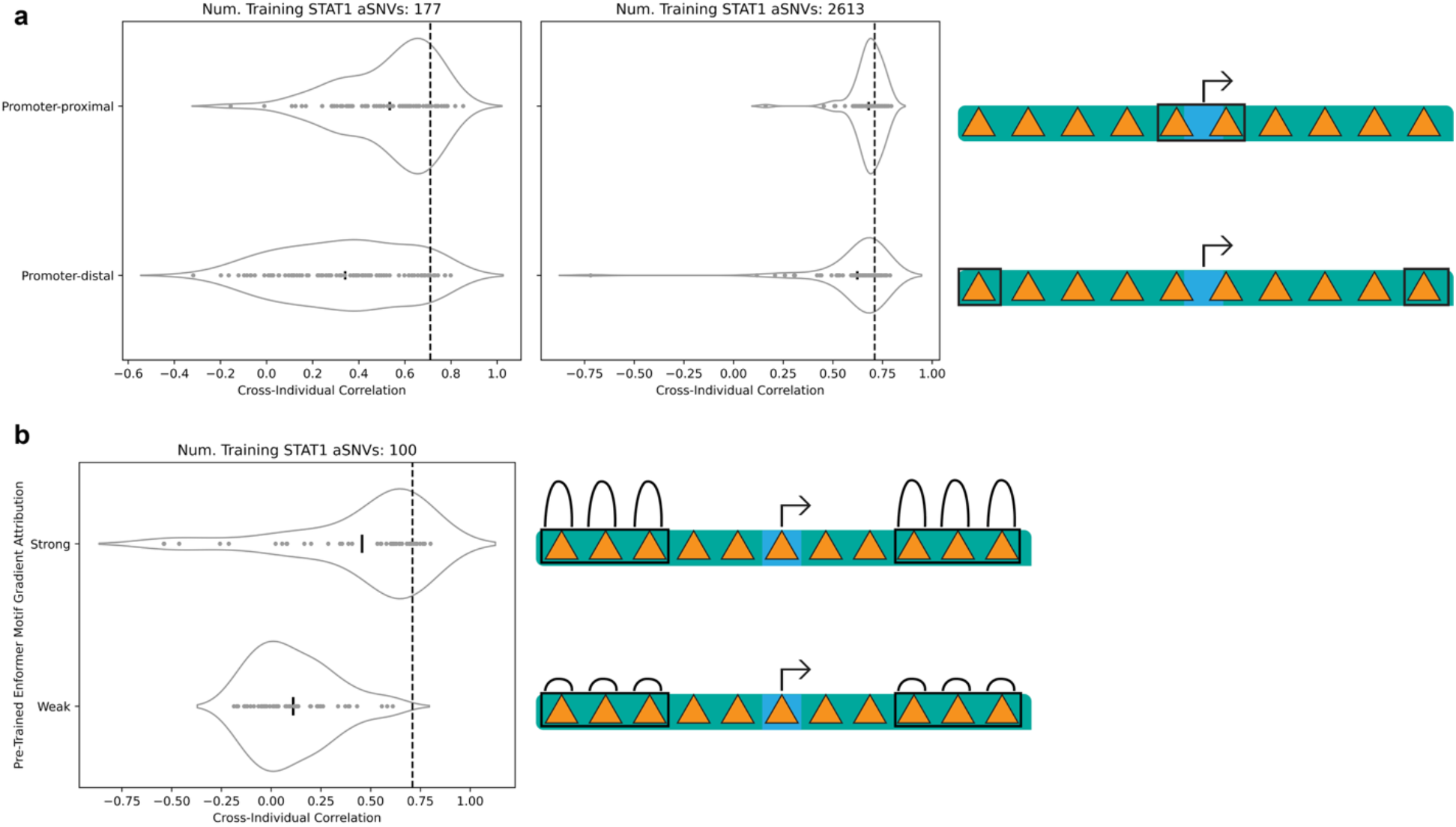
Miscalibrated pre-trained priors cause enhancer-to-gene linking issues that hinder learning of variant effects. **a)** Cross-individual correlation results for 87 test set genes (HOGP, **Methods**) where aSNVs are placed either within 6kb of the TSS (Promoter-proximal) or within 6kb of either the upstream or downstream window edge (Promoter-distal) during training and evaluation. These models were trained on either 177 aSNVs (left) or 2613 (right). Enformer models learn and generalize promoter-proximal variant effects better than distal ones, implying issues linking distal elements housing these variants to their target genes, a gap that is partially rescued with increased data. **b)** Similar to (**a**), except the two models are trained on aSNVs that perturb motifs that are not Promoter-proximal (are at least 6kb away from each gene’s TSS), are distance matched between the two groups, and instead differ primarily by whether pre-trained Enformer originally had strong or weak gradient attributions around these motifs (**Methods**). Evaluations are on 50 test set genes (HOGP, **Methods**). During post-training, Enformer more readily learns distal variant effects from distal regions that already have high importance.

Consistent with the literature^11–13,16–19^, we used elastic net models trained directly on genes from the held-out set, and evaluated on the same genes but held-out people (HOP), to generate an expected result a model could reasonably achieve without needing to learn their effects from elsewhere in the genome (**Methods**). While average HOGP accuracy improved with increased training dataset size, it consistently underperformed the elastic net baseline, even after increasing the dataset size to include ∼12,000 genes (with one aSNV per gene), the maximum number of Whole Blood-expressed protein-coding genes in Enformer’s original train set (**Fig. 2a**). No matter the training dataset size, this underperformance was inconsistent between genes, with HOGP accuracy appearing high for some genes but not others.

To investigate the causes of this underperformance, we first analyzed the predicted effect sizes of aSNVs relative to surrounding naturally observed SNVs. We previously showed that such underperformance on the personal genome task stems from underestimation of causal variant effects relative to nearby variants with weaker or no effects, and that this underperformance is therefore a proxy for unsuccessful fine-mapping^10^. Consistent with this, aSNV predicted effect sizes were much weaker for less performant genes, and often had near-zero predicted effect sizes that were even weaker than many other naturally occurring SNVs found in the same locus – despite the outsized expression effect the aSNV was designed to have relative to these natural SNVs (**Supp. Fig. 1a-b**) – but stood out prominently above these other SNVs for more performant genes that matched the elastic net accuracy (**Fig. 2d**).

Additionally, to ensure this personal genome prediction underperformance could not be explained instead by inaccurate attributions for naturally occurring SNVs also found in these individuals, we fit these elastic net baselines two ways: using all SNVs (including the aSNV) in the region, and using the aSNV as the only covariate (**Methods**). The former only slightly outperformed the latter by capturing this additional natural variation, consistent with the aSNV’s outsized importance to individual gene expression differences relative to all other SNVs in a given gene window, and that attainable accuracy in these loci depends almost entirely on its prominent detection (**Supp. Fig. 1b**). As fine-tuned Enformer underperformed even the more lenient baseline that considers the aSNV alone, this underperformance cannot be explained by inaccurate natural variant effect attributions, and our results suggest it is instead caused by underestimation of aSNV effect sizes. Elsewhere in this manuscript, we therefore use the aSNV-only baseline to ask only that fine-tuned Enformer explain the gene expression differences the aSNV alone confers. We interpret underperformance relative to this baseline as a proxy of incomplete fine-mapping of the aSNV, where the aSNV is not weighted with sufficient strength relative to surrounding background SNVs in a manner that is consistent with its true outsized importance to expression, consistent with our past findings^10^.

For a subset of 100 training genes, we set the aSNVs’ MAF to 0% for individuals in the train set, and reintroduced them at 50% MAF in HOP as a proxy for rare genetic variants that are not observed around a locus during training but appear in other individuals during test time. We found personal genome prediction accuracy from individuals in the validation set, where the aSNV is being observed around these 100 train genes for the first time by Enformer, plateaued around the same time as totally held-out genes (HOGP; **Supp. Fig. 4**). These results indicate S2F models learn to generalize held-out variant effects for loci seen during training with similar efficiency to those from completely unseen loci. This suggests post-training efforts to improve S2F rare variant predictions face similar challenges to variants from totally unseen loci, and we focus primarily on the latter throughout the rest of this manuscript.

**Figure 4:**
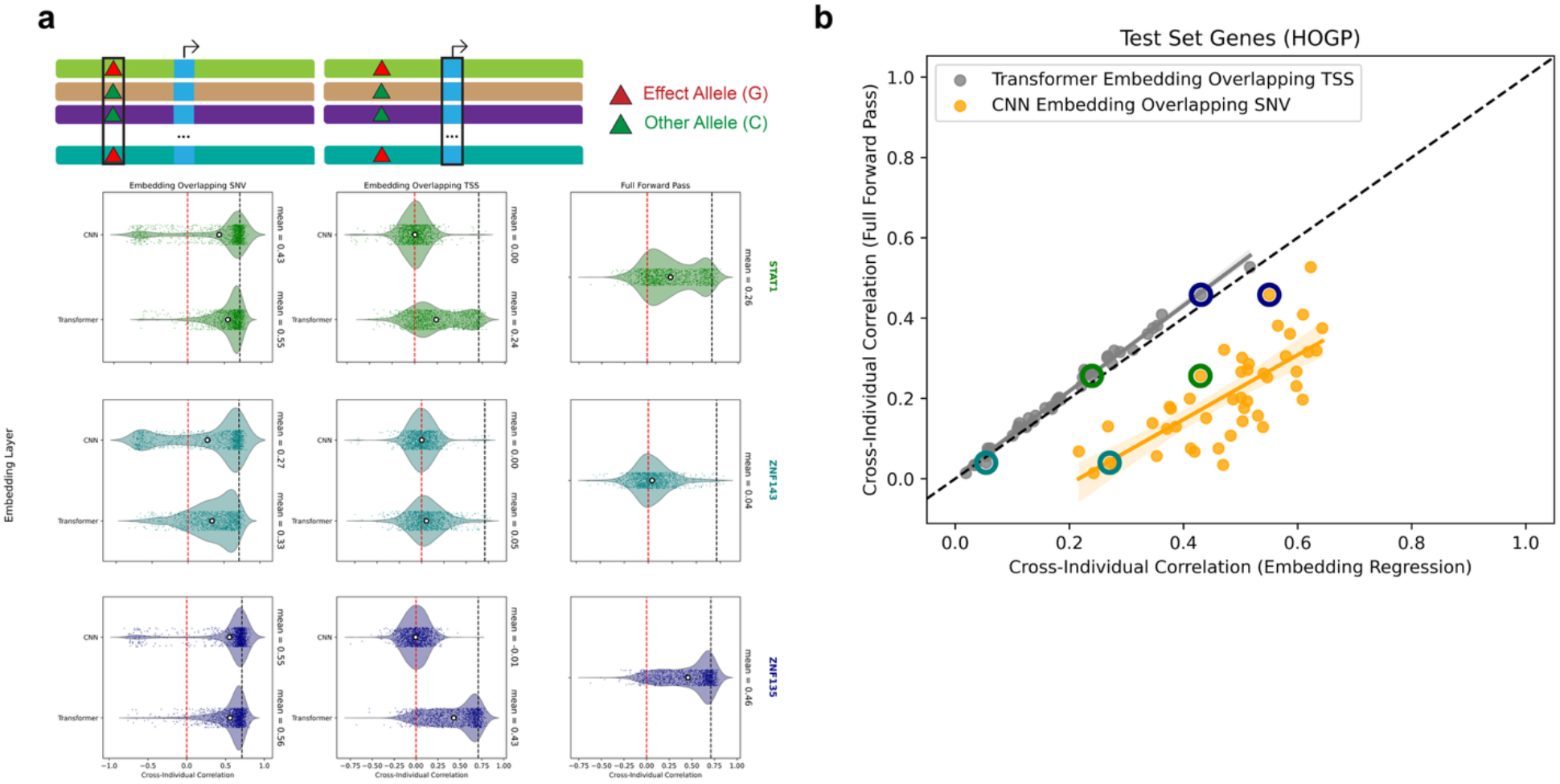
Demonstration of incomplete enhancer-to-gene linking hindering learning of aSNV effects. **a)** Cross-individual correlation results from 1,183 test set genes (HOGP) after fitting linear models to predict gene expression from final layer CNN or Transformer embeddings (rows within each panel) from the position along the sequence axis containing the aSNV (wherever it may be for a given gene, left column) or the center position corresponding to the TSS (right column). These embeddings come from fine-tuned Enformer models trained using LAMP. Fine-tuned Enformer predictions (not from models fit on embeddings) are shown in the right column. Results from models trained on aSNVs perturbing one of 3 TF motifs (STAT1, ZNF143, ZNF135) is shown. Embeddings overlapping the aSNV position are relatively informative for predicting individual gene expression, outperforming predictions that come from the full forward pass, but this information is incompletely propagated to the TSS via attention, leading to less accurate predictions from linear models fit on the TSS Transformer embeddings. This incomplete propagation of information reflects incomplete E2G linking. TSS CNN embeddings are a negative control. **b**) Each point represents the mean HOGP cross-individual correlation for one of 45 fine-tuned Enformer models trained and evaluated on aSNVs perturbing motif instances for one of 45 different TFs. X-axis represents predictions coming from regression models fit on CNN aSNV embeddings (orange) or TSS Transformer embeddings (grey). Two points along the horizontal axis therefore represent results from different embeddings and sequence positions but the same motif. Y-axis represents predictions coming from the fine-tuned Enformer full forward pass, rather than regression from its embeddings. A similar amount of genotype information is always lost between CNN aSNV embeddings and TSS Transformer embeddings (gap between orange and grey along the horizontal axis), no matter the motif, reflecting similar E2G linking deficits after fine-tuning on aSNVs perturbing any motif. Predictions deriving from TSS Transformer embeddings correlate highly with those from full forward passes, indicating the extent to which aSNV genotype information is propagated from its position to the TSS determines predictive accuracy. Motifs for which HOGP predictions are less accurate are also associated with less informative CNN aSNV embeddings, suggesting confident detection of the aSNV by the CNN module also presents a learning obstacle (an observation that is further explored in Fig. 5). Points with a colored border represent TF motifs from (**a**).

Personal genome prediction underperformance held after modifying a variety of hyperparameter choices (**Supp. Figs. 5a-b**), as well as replacing Enformer with Borzoi (**Supp. Figs. 5c-d**). We also attempted to control for various confounders from our setup, including the presence of natural genetic variation in STAT1 motifs and the natural presence of the G allele (“effect allele”) at the aSNV’s relative position in other STAT1 motifs that were not perturbed by the aSNV, concluding these only minimally impacted results (**Supp. Fig. 6**). In addition, we altered the training setup to train directly on individual aSNVs and natural SNVs, rather than combined into personal genome sequences, finding that this too minimally altered results (**Supp. Fig. 7**). Finally, we systematically varied the aSNVs effect size and MAF, finding that although decreased MAF hindered learning of variant effects, none of these changes rescued HOGP accuracy relative to the elastic net baseline (**Supp. Fig. 8)**. These results suggest it is unlikely that this underperformance can be explained by hyperparameter choices or other obvious modeling and architectural decisions nor can it be caused by the difficulty in learning from the complexity of real genetic variation due to the idealized causality we simulated using LAMP. Instead, we hypothesized that underperformance reflects a fundamental challenge in post-training S2F models to learn how a variant’s motif perturbation affects expression and generalize it to new loci, consistent with past failures fine-tuning with unaltered personal genome datasets. Given widespread interest in post-training S2F models, including with genetic variation, personal genome, perturbational, or evolutionary datasets, this motivated us to further investigate.

**Figure 5:**
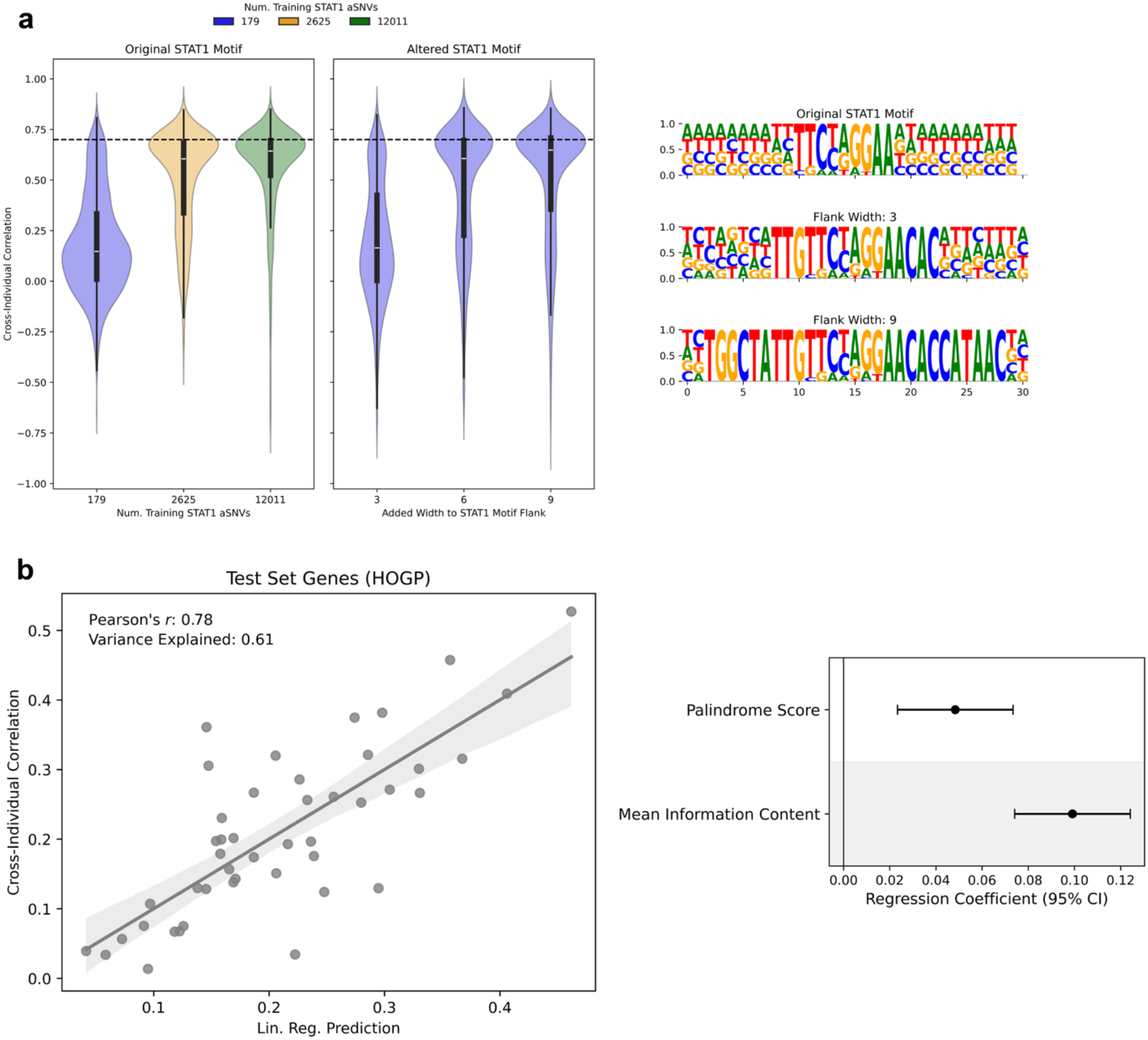
Sequence context variability obscures learnable patterns of variant-induced motif perturbations during post-training. **a)** HOGP performance on 88 test set genes after adding additional nucleotides to the flanks of STAT1 motifs and perturbing 179 STAT1 motifs with aSNVs (right violin plot). All candidate STAT1 motifs (**Methods**) are altered in this way, including those not perturbed by the aSNV. Logo plots on the right show the average nucleotide composition at positions relative to the motif before (top) and after adding 3 or 9 nucleotides to the flank. Violin plots on the right show results before modifying STAT1 motifs in this way, and are matched in experimental design (aSNV genomic positions, selected genes and individuals, etc.) except for the STAT1 flanking sequence modification and two models are trained on more than 179 aSNVs. By only introducing additional, fixed flanking nucleotides to the STAT1 motif, and thereby decreasing sequence context variability around the aSNVs, Enformer more efficiently learns the motif perturbation pattern, performing comparably to the 12,011 aSNV model fine-tuned with unaltered STAT1 motifs. **b**) Average HOGP performance on 1,183 test set genes after fine-tuning 45 Enformer models on aSNVs perturbing one of 45 different motifs (same experiment as Fig. 4a**-b**) is displayed on the Y axis. Each point is the HOGP cross-individual correlation from one of these models, averaged over the test set genes. X-axis shows predictions from a linear regression model fit to predict y-axis HOGP values after being fit using only each motif’s palindrome score and length-controlled IC as covariates (**Methods**). Increased palindromicity and IC amount to decreased sequence context variability, and these two motif features have a significant positive association (*p* = 3.4 x 10^-4^ and *p* = 5.7 x 10^-10^, respectively; right panel) with HOGP accuracy, and fit a linear model that robustly explains differences in HOGP after fine-tuning using aSNVs that perturb different motifs (left panel). These data suggest sequence context variability impacts post-training of variant effects.

**Figure 6:**
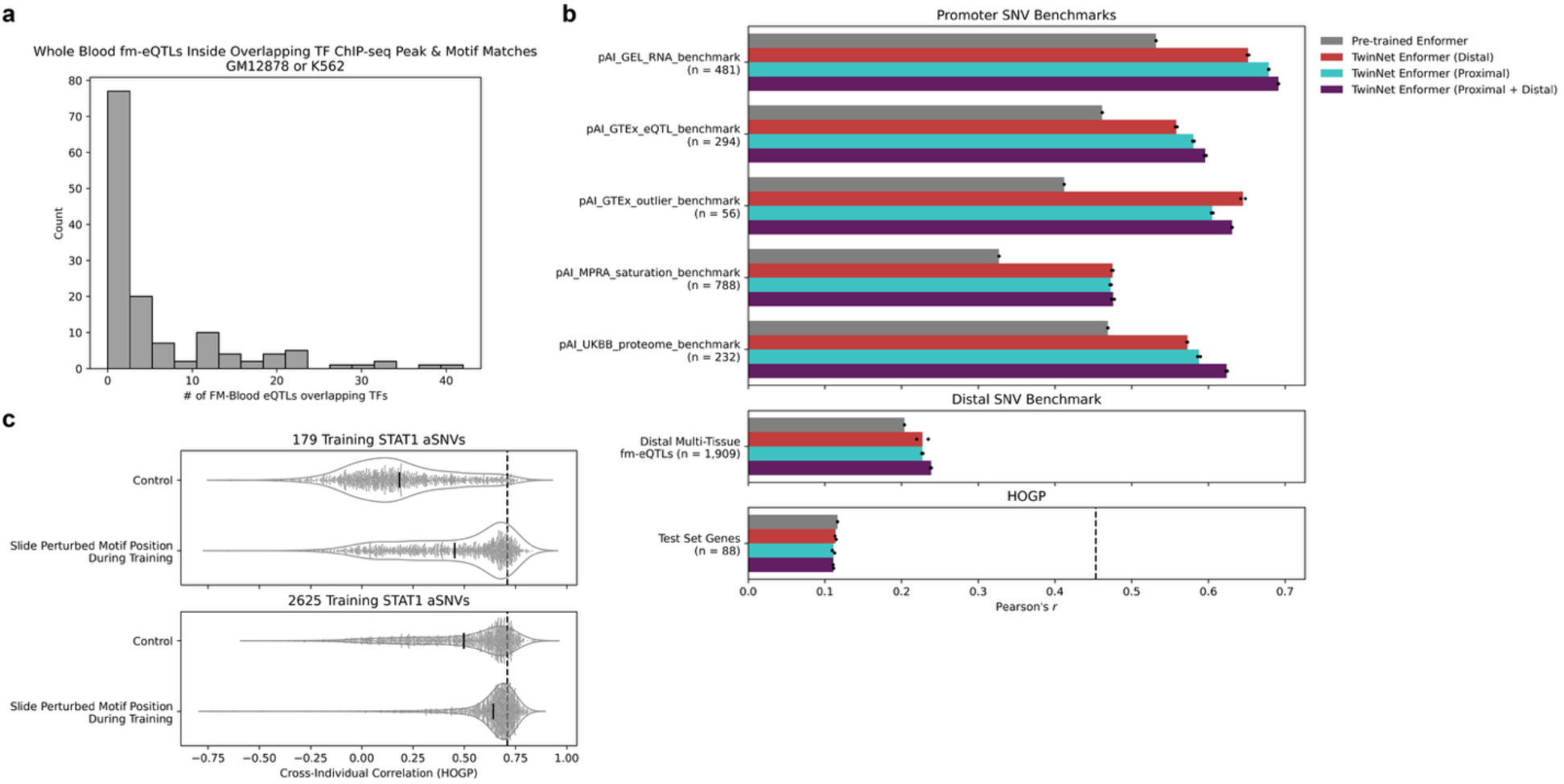
LAMP guides the design of post-training datasets beyond natural genetic variation. **a)** The number of Whole Blood fm-eQTLs found within both a TFs motif and its ChIP-seq peak in blood-relevant (K562, GM12878) cell lines. Most TFs have 0 or 1 fm-eQTLs in a binding site genome-wide, whereas thousands of aSNVs perturbing different instances of the same TFs same binding site were needed to optimize learning of its variant effects (Fig. 2). While the rest of this manuscript focuses on the ∼49kb window around the TSS, we included fm-eQTLs that are any distance away from their target gene. **b**) TwinNet Enformer models fine-tuned on rare expression outlier SNVs (**Methods**) form improved promoter SNV predictions (top), but gains are negligible for distal SNVs (middle) and personal genome predictions (bottom), which depend on improved distal SNV predictions^10^. Promoter SNV benchmarks come from the promoterAI (pAI) manuscript^15^. **c**) Cross-individual correlation (HOGP) after implementing a data augmentation strategy that repositions aSNV-perturbed STAT1 motifs to break pre-trained priors regarding distance leads to more data efficient learning of aSNV effects. Controls are the same models used in Fig. 2a. Each point is a gene from the held-out test set.

### Strong, miscalibrated pre-trained priors impair linking of distal enhancers to target genes, hindering learning of variant effects

We first hypothesized that the per-gene inconsistency in HOGP underperformance (**Fig. 2a**) reflects, at least in part, differential sensitivity to STAT1 motifs based on their distance from the target gene, consistent with the well-documented neglect of distal regulatory elements by S2F models^1–3,14^. Because aSNVs were placed in randomly chosen STAT1 motifs, which are uniformly distributed across the 49kb window (**Supp. Fig. 2a**), they fall near the promoter for some genes and far from it for others. To directly investigate this, we placed STAT1 aSNVs into personal genome sequences in positions either near (+/- 6kb of TSS) or far (within 6kb of either window edge) from the TSS of train and held-out genes, holding the genes, individuals, and target expression values identical between the two groups so that aSNV position was the primary difference (**Methods**). As hypothesized, we found that fine-tuned Enformer preferentially learned and generalized the STAT1 aSNV effects from promoter-adjacent regions, and struggled with aSNVs in distal motifs, a deficit that was reduced but persisted with a large increase in training dataset size (**Fig. 3a**). We repeated this experiment by selecting only STAT1 motifs within ENCODE4^23^ candidate cis-regulatory elements (cCREs) to perturb with aSNVs, obtaining nearly identical results, indicating Enformer’s insensitivity to distal aSNVs persists even among the distal motifs most likely to be within genuine functional elements (**Supp. Fig. 9a**). Therefore, unless otherwise stated, we did not restrict to cCRE motifs elsewhere, as doing so reduces the number of eligible motifs per gene and limits our ability to match other experimental parameters across conditions. These results suggest S2F models struggle to learn distal variant effects, implying challenges learning to link enhancers and other distal regulatory elements housing these variants with target genes (enhancer-to-gene linking; E2G linking).

We next hypothesized that weak E2G linking is not a gap Enformer learned during fine-tuning, but instead reflects strong, well-documented pre-trained priors^1–3,14^ against distal regulatory elements by Enformer and other S2F models. We speculated that promoter bias in pre-trained S2F models leads to substantially more data being required during fine-tuning in order to successfully strengthen distal E2G links. To investigate, we computed the gradient of Enformer’s gene expression predictions with respect to each input DNA sequence, measured before fine-tuning (**Methods**). This value reflects the importance of each nucleotide to the gene expression prediction, so we used it as a proxy of the strength of pre-existing E2G links. We then placed STAT1 aSNVs into distal STAT1 motifs (beyond +/- 6kb of TSS) with either weak or strong pre-existing attributions, matching the two groups for distance from the gene’s TSS such that attribution strength was the primary difference (**Methods**). Consistent with our hypothesis, we found that fine-tuned Enformer generalized the STAT1 aSNV effects significantly better to HOGP when they were placed in distal regions it already attended to strongly before fine-tuning, relative to distance-matched regions with weaker pre-existing attributions (**Fig. 3b**). Repeating this experiment after placing aSNVs in motifs within cCREs again led to very similar results (**Supp. Fig. 9b**). These findings suggest Enformer learns a new variant effect quite readily within distal regions it already considers important, but struggles from distal regions with weak pre-existing attributions. Most distal regions receive weak attributions (**Supp. Fig. 10a**), however, including many of those containing fine-mapped eQTLs (fm-eQTLs; **Supp. Fig. 10b**) and those with experimentally verified regulatory importance^14^. Together, these data suggest Enformer inefficiently strengthens E2G links during post-training to regions it learned to downweight after pre-training, and this leads to inefficient learning of distal variant effects because distal regions are particularly prone to downweighting. We speculate that strong miscalibrated pre-trained priors regarding distance, and the resulting data inefficiency in learning from distal variants, contribute to the failure of previous attempts to fine-tune S2F models on personal genomes.

To directly demonstrate that incomplete E2G linking hinders learning of variant effects, we analyzed fine-tuned Enformer’s HOGP embeddings, after placing aSNVs in one randomly sampled STAT1 motif per gene, from two positions in the input sequence — the bin overlapping each STAT1 aSNV, wherever it sits for a given gene, and the bin containing that gene’s TSS — taken at both the final convolutional and final transformer layers, for four embedding types in total. For each of the four embedding types, we fit a regression model to predict individual gene expression values using embeddings from Enformer’s train genes and individuals, and we evaluated it on embeddings of the same type from the HOGP set (**Methods**). This framework measures the extent to which each type of embedding captures the relationship between aSNV genotype and gene expression in HOGP. Regressions on aSNV-overlapping embeddings measure how well the aSNV is detected locally, regressions on TSS-overlapping embeddings measure how much of that detected signal has been propagated over distance to the gene’s TSS, and aSNV genotype information lost in between these steps reflects incomplete E2G linking. Convolutional and transformer embeddings from bins overlapping STAT1 aSNVs supported HOGP predictions that were much more accurate than that of the full forward pass, and only modestly underperformed the elastic net baseline (**Fig. 4a**), indicating genotype information is largely preserved at the aSNV position throughout the model. Embeddings from the TSS bin behaved differently. Convolutional embeddings from this position were not informative for predicting individual expression differences, as expected given that information from the STAT1 aSNV sequence position has not yet propagated to the TSS at this depth in the model, and is only expected to occur after the transformer module. Indeed, transformer embeddings from the TSS bin were more informative and performed comparably to the full forward pass, yet remained much less informative than embeddings taken from the aSNV position itself (**Fig. 4a**). These data suggest aSNV genotype information is incompletely propagated from its position within the sequence to the gene TSS via attention and this loss of information, representing incomplete E2G linking, weakens variant effect predictions. In other words, the transformer is not fulfilling its intended purpose. Rewiring attention weights to propagate aSNV genotype information to the target gene is therefore an important limiting step during post-training, and our earlier results suggest this rewiring is data inefficient because pre-trained priors bias Enformer from attending to distal regions.

We repeated this experiment, replacing STAT1 with the motif of one of 45 other Whole Blood-expressed transcription factors (**Methods**), resulting in 45 fine-tuned Enformer models. As motifs differ in length and information content (IC) at each position, we chose to place aSNVs in the highest IC position of each motif rather than a fixed position, including for STAT1, whose aSNV here and in the preceding embedding analysis therefore sits at a different position in the motif than in our earlier experiments (**Methods**). This was done to control for a weak relationship between the IC at a motif’s position and HOGP accuracy (**Supp. Fig. 11**). We found HOGP accuracy from each model’s full forward pass is tightly correlated with the extent to which aSNV genotype information has been propagated to the TSS overlapping transformer embeddings, reflecting the importance of E2G linking for variant effect predictions (**Fig. 4b**, grey). In every case, embeddings overlapping aSNVs are more informative for explaining gene expression differences, and therefore better capture aSNV genotype information, than transformer embeddings overlapping the TSS. Thus, for every motif, we observed a substantial loss of aSNV genotype information between embeddings overlapping the aSNV versus those overlapping the TSS (**Fig. 4b**, orange vs. grey), further suggesting that incomplete E2G linking hinders accurate learning of variant-induced gene expression effects.

### Sequence context variability obscures learnable patterns of variant-induced motif perturbations during post-training

We next investigated whether, independent of E2G linking challenges, local detection of the aSNV’s motif perturbation acts as an upstream bottleneck. We found that the informativeness of aSNV-overlapping CNN embeddings was associated with both the informativeness of the TSS-overlapping transformer embeddings and the accuracy of HOGP predictions from the full forward pass (**Fig. 4b**, orange vs. grey). This is exemplified by ZNF143 (lower informativeness) and ZNF135 (higher informativeness; **Fig. 4a**). This suggests weaker detection of the aSNV in early convolutional layers leaves less genotype information available to propagate, and that this weaker information is then further lost after the transformer, yielding poor E2G linking and worse HOGP predictions. This is consistent with our past observations, where failure by AlphaGenome to detect local variant effects onto enhancer activity hampers variant effect predictions, independent of distance^10^. While the loss of aSNV genotype information during E2G linking was highly consistent between TFs (**Fig. 4b**, gap between orange and grey along horizontal axis), local detection issues were inconsistent across TFs (**Fig. 4b**, spread between orange points along x-axis).

As the choice of TF motif was the primary difference between these experiments, we hypothesized that the motif sequences themselves account for the differences in local detection. More broadly, we hypothesized that the aSNV motif perturbation pattern becomes easier to learn, and distinguish from the same nucleotide substitutions occurring elsewhere, when the sequence surrounding the aSNV differs less between examples. Properties of motifs housing the aSNV that reduce this sequence context variability — greater length, higher IC, and palindromicity — would theoretically make the aSNV easier to detect by causing the same nucleotides to recur around it across loci and DNA strands, rather than a comparatively variable set. To test this directly, we artificially extended all significant STAT1 motif matches in the genome before introducing one aSNV per gene, fixing more of the sequence surrounding the aSNV and thereby reducing sequence context variability between examples (**Methods**) by increasing motif length and average IC. HOGP accuracy was highest with longer flanks, approaching the elastic net baseline (**Fig. 5a**). Under these conditions, a model trained on 179 aSNVs performed comparably to one trained on approximately 12,000 (median HOGP *r* = 0.648 vs. 0.643), despite the only additional difference in train/test data being the artificial flank added to all STAT1 motif instances.

Consistent with this, after fine-tuning Enformer on aSNVs perturbing each of 45 motifs, we found each motif’s palindromicity and length-controlled IC had a significant positive effect (*p* = 3.4 x 10^-4^ and *p* = 5.7 x 10^-10^, respectively, from linear regression) on HOGP performance, and these two features were highly informative in predicting HOGP performance after training on each motif, explaining 61% of this variance (**Fig. 5b**, Pearson’s *r* = 0.78). Thus, motif properties that decrease sequence context variability are associated with more data-efficient learning of aSNV effects, and this helps explain the discrepancy in aSNV detection between different motifs from our embedding analysis (**Figs. 4a-b; Supp. Fig. 12**) as well HOGP accuracy differences when perturbing different motif positions with the aSNV (**Supp. Fig. 11**). As shorter motifs are more likely to appear in the genome by chance, we verified this trend was independent of the prevalence of each of these motifs in the genome (**Supp. Fig. 13**). We also verified its independence with each TF’s Whole Blood expression level and with Enformer’s pre-trained gradient attributions around each motif, which we observed to be an important indicator (**Fig. 3b**). Together, these results suggest that the extent to which the sequence surrounding an aSNV varies between loci — greatest for short, low IC, and less palindromic motifs — obscures the motif perturbation pattern caused by the variant and makes it harder to learn. Sequence context variability may therefore be an underappreciated genomic confounder when post-training S2F models on genetic variation.

### Observational genetic variation datasets are likely insufficient to optimize S2F variant effect predictions

We next asked whether observed human genetic variation contains enough strong, common, causal variants within TF motifs to match the number of training aSNVs (also strong, common, and causal) we needed to optimize HOGP predictions. To this end, we counted, for each TF, the number of Whole Blood fm-eQTLs (posterior inclusion probability > 0.9) overlapping any of its ENCODE4 GM12878 and/or K562 ChIP-seq binding sites genome-wide (Methods), mirroring our aSNV setup, in which every aSNV perturbs a binding site for the same TF. Each TF’s binding sites overlapped a median of 2 fm-eQTLs, with most overlapping 0 or 1, and no TF’s binding sites overlapping more than 42 (**Fig. 6a**). By contrast, our LAMP analyses introduced aSNVs into up to 12,011 STAT1 binding sites (**Fig. 2a**), which was still insufficient to optimize HOGP accuracy. Additionally, aSNVs were designed to exhibit directionally consistent effects, the same outsized effect size for each gene, perturb the same relative position of the motif with the same two alleles, and appear uncorrelated with the presence of other observed variants to simplify the learning task; natural SNVs do not. Besides fm-eQTLs, rarer or weaker causal variants also exist, but modeling aSNVs at lower MAFs or with weaker effect sizes likewise led to less efficient learning (**Supp. Fig. 8**), so their consideration is unlikely to resolve this shortfall. Strong, common, naturally observed causal variants are therefore likely to be both less abundant and harder to learn from than aSNVs. We conclude that the scale of observed human genetic variation is vastly insufficient to maximize accuracy of HOGP variant effect predictions via simple fine-tuning.

### Using LAMP to guide future experimental design

As a proof of concept for using LAMP to optimize and troubleshoot S2F post-training design, we attempted to corroborate our finding that S2F models are better primed to learn variant expression effects from within promoters versus distal regions (**Figs 3a-b**). We used a ‘TwinNet’ contrastive architecture to fine-tune Enformer to predict the effect size of rare multi-tissue expression outlier variants from GTEx while passing in the two alleles of each variant as contrastive training examples (**Methods**), differing from our previous personal genome training setup. A similar setup was used to fine-tune promoterAI^15^, where it demonstrated improved predictive accuracy on promoter expression-modifying variants. Consistent with this, fine-tuning Enformer on promoter and/or distal expression outlier SNVs led to strong improvements in promoter variant effect predictions from a variety of benchmarks (**Fig. 6b**, top; **Supp. Fig. 14a**). Shuffling the effect sizes associated with each training SNV led to stochastic outcomes that did not reliably beat pre-trained Enformer, indicating fine-tuning on this curated set of SNVs was necessary for the improvement (**Supp. Figs. 14a-b**). By contrast, fine-tuning on promoter outliers (n = 5,447 SNVs), distal outliers (n = 122,293 SNVs), or both led to minimal improvement on distal fm-eQTLs, consistent with our finding that pre-trained priors leave distal E2G links data inefficient to strengthen (**Fig. 6b**, middle). Consistent with our past finding that improving personal genome predictions depends largely on better fine-mapping of distal variant effects^10^, we also find these models did not form improved gene expression predictions from personal genome sequences (**Fig. 6b**, bottom; **Methods**). This is an important negative result that addresses a previously open question: while fine-tuning on rare promoter outliers improves predictions for unseen promoter variants, the same setup does not yield gains for distal variants. But improving S2F personal genome predictions and fine-mapping capabilities depend upon better distal variant predictions (**Figs. 3a-b**). This demonstrates the value of the LAMP framework, which anticipated this outcome, and offered an avenue into understanding its cause.

We therefore used LAMP to test a future experimental design strategy. Inspired by past genomic augmentation strategies^24^, we sought to promote invariance to the position of the STAT1 motif within the input sequence, with the goal of promoting E2G link formation to distal regions and more data-efficient learning of distal variant effects. This was intended to counteract our finding that strong, miscalibrated pre-trained priors bias S2F models away from linking distal elements to target genes, and in turn away from learning the effects of variants within those elements (**Figs. 3a-b**). We modified the LAMP training scheme by sliding aSNV-perturbed STAT1 motifs from their endogenous position elsewhere during training, while keeping gene expression targets the same (**Methods**). This form of augmentation is meant to break pre-trained priors regarding distance while keeping the number of aSNVs and genes for training the same, rather than by increasing the amount of training data – which is limited in real scenarios. We found this led to significantly improved performance on HOGP (**Fig. 6c**). Underlying this improvement was an overall increase in gradient attributions around STAT1 motifs, as well as a weaker dropoff in these attributions around STAT1 motifs that sit further from the TSS (**Supp. Fig. 15**). Looking forward, we envision a similar strategy can be applied by artificially moving distal regulatory elements throughout the sequence during post-training to break miscalibrated pre-trained priors regarding distance. To prevent overfitting to the positional invariance introduced through augmentation that may not align with real data, a final fine-tuning step with unaltered data can be employed, as others have recommended^24^. Additionally, motif perturbation patterns for some motifs may be harder to learn than others (**Fig. 5b**), and these motifs may need to be oversampled in post-training datasets; LAMP could be used to identify these motifs and optimize their sampling.

## Discussion

We introduced LAMP, a framework for troubleshooting and optimizing S2F post-training objectives via tracking how S2F models learn aSNV effects. Through a series of *in silico* experiments using Enformer, we showed that S2F models are poorly calibrated for predicting the effects of motif-disrupting distal variants on gene expression and data inefficient for learning these effects during post-training. Because causal variant effects in human observational genetic variation datasets are likely harder to learn from than aSNVs— being likely less abundant, less stereotyped in their motif perturbations, and confounded by LD — it is unlikely that such datasets will be sufficient to optimize personal genome predictions, and therefore fine-mapping objectives, by S2F models through simple fine-tuning.

Critically, LAMP not only illuminated this data inefficiency, but helped explain the underlying sources. This framework identified two major barriers to learning and generalizing variant effects during post-training: (1) Challenges learning to strengthen E2G links to distal regulatory elements containing causal variants, caused by strong, miscalibrated pre-trained priors that bias attention weights to neglect distal regulatory elements; and (2) natural sequence context variability acting as an underappreciated genomic confounder by obscuring these motif perturbation patterns, further hindering the learning of distal variant effects from observational genetic variation datasets. We speculate the latter issue is only more pronounced for natural (non-aSNV) variants, whose effects on regulatory sequence are far less stereotyped than the aSNVs studied here. We note that S2F models form strong local accessibility and TF binding predictions^2,3^, including across individuals^13^ and for lower-affinity motifs^25^. Our results regarding challenge (2) are not in conflict with these findings, as they correspond only to the extent to which sequence context variability obscures motif perturbation patterns by genetic variants during post-training on the gene expression prediction task, independent of the ability of S2F models to learn the importance of these motifs for local readouts during pre-training.

Generalizing beyond aSNVs to real data, these obstacles explain why S2F models continue to underestimate causal variants relative to their neighbors^10^ — a major S2F challenge underlying poor personal genome prediction and fine-mapping — after post-training on human genetic variation datasets^16–19^. Bridging this gap will likely require perturbational datasets that disproportionately sample distal elements and sample each motif deeply enough to overcome sequence context variability, data augmentation strategies that also target these issues, or both, and we propose LAMP as a tool for optimizing such experimental designs. For example, our results suggest that data augmentation strategies that explicitly target challenge (1), by promoting invariance of the aSNV effect to its distance from the TSS, lead to more data-efficient learning of aSNV effects. Others have proposed similar augmentation strategies, followed by fine-tuning on unaugmented data to prevent overfitting to the augmentations^24^; our results further suggest such strategies may improve learning of variant effects.

LAMP has several limitations. First, fine-tuning on personal genomes and paired expression values is highly compute intensive. Second, we focus on Enformer, rather than newer generation S2F models like Borzoi and AlphaGenome. While these models reportedly outperform Enformer on variant effect and personal genome tasks, both still fall short of expectation on the personal genome task and retain miscalibrated priors that bias them away from distal elements, so we expect our findings to hold for them as well^2,3,11–14^. Indeed, we observed similar results for a subset of experiments with Borzoi, while not investigating AlphaGenome with LAMP due to the size of the model. Third, our approach simulates learning of variant gene expression effects through a motif perturbation mechanism, and does not apply to other types of variants that act through different mechanisms. Fourth, this approach only considers aSNVs perturbing one TFs motif at a time. In reality, models learning from natural variation may draw on regulatory logic shared between motifs — for instance among members of the same TF family — an advantage our single-motif setup does not capture. Lastly, because LAMP simulates variant effects, learning from real genetic variation is not guaranteed to behave identically. Still, many of our main findings agree closely with the field’s broader understanding of E2G linking issues in S2F models, which supports their relevance to learning real variant effects during post-training.

## Methods

### Personal Genome Sequences

We generated personal genome sequences for individuals from GTEx v8 as part of a different study^16^. Briefly, we used *bcftools consensus* to replace reference genome (hg38) nucleotides with SNVs present in each individual’s unphased whole genome sequencing (WGS) VCF. We generated two of these consensus files per individual, to handle cases of heterozygosity. We one-hot encoded each, and averaged them together to generate a single input for model training and inference. This average encoding reflects whether each individual is homozygous for the reference allele (0/0; encoded as a 0), heterozygous (0/1; encoded as 0.5), or homozygous for the alternate allele (1/1; encoded as a 1) for each observed SNV.

### Gene Expression Data

Starting with gene read counts (GTEx_Analysis_2017-06-05_v8_RNASeQCv1.1.9_gene_reads.gct.gz) from individuals with both RNA-seq and WGS data in at least one of 49 GTEx tissues, we computed TMM normalization factors across all tissues, using a python port of edgeR^26^ (https://github.com/broadinstitute/pyqtl/blob/master/qtl/norm.py). We then converted these counts to a Counts Per Million (CPM) matrix after applying these factors and then log transformed the expression values. Following direction from GTEx, we kept genes with at least 6 reads in at least 20% of samples (we refer to these genes as “expressed” in Whole Blood throughout the rest of this manuscript). We implemented this procedure to normalize expression values across tissues to enable future analyses that compare expression across tissues. We used this gene expression matrix as the starting point for the gene expression values used elsewhere in this manuscript (see **Targets** for more information).

### Implementation of LAMP: Fine-tuning Enformer on aSNVs embedded into personal genome datasets

#### Inputs

We started with personal genome sequences created in **Personal Genome Sequences**. Unless otherwise stated, we trained on sequences of length 49,152 base pairs (bp) centered on the target gene’s TSS (as annotated by GENCODE^27^ v26). For each gene, we pick one motif to perturb with the aSNV (see **Selection of Motifs to Perturb with aSNV** for more detail), and we randomly assign each individual to receive either the “effect allele”, whose presence will correspond to an increase in gene expression (see **Targets**), or the “other allele”, whose presence will correspond to no change in gene expression.

We define the aSNV to exist in the same (strand-aware) relative position of every perturbed motif in each experiment, and the effect allele is defined as the nucleotide with the highest JASPAR^22^ PWM value at that position, modeling the scenario where increased binding affinity causes an increase to gene expression. The other allele is defined as the nucleotide with the lowest PWM value at that position. For the STAT1 aSNV placed at position 6, that corresponds to a G “effect” allele and a C “other” allele when the motif is on the (+) strand. Each individual receives only homozygous dosage of their assigned aSNV genotype; we did not introduce aSNVs as heterozygous SNVs.

During training, we look up the assigned aSNV genotype for each individual in the batch, and substitute their randomly assigned allele into the corresponding position within the motif on the fly using tangermeme^28^. If one individual is assigned the effect allele for the aSNV around one gene, they may receive the other allele for a different gene. If the chosen motif is on the (-) strand, the motif appears as its reverse complement when (+) orientation sequences are passed in. To alter the same relative position of the binding site, we therefore reverse complemented the aSNV’s appearance within these motifs (e.g., if the aSNV is meant to be placed at the first position within the motif as an A/C, it is instead placed at the last position as a G/T). This causes the aSNV to alter the same motif position, no matter which strand it is on. We randomly reverse complement during training so the model learns both orientations.

### Targets

We started with the gene expression matrix created in **Gene Expression Data**, focusing on the Whole Blood GTEx tissue. We aimed to modify each individual’s gene expression values for each gene to embed into the data a strong causal relationship between the presence of the aSNV effect allele and a consistent increase in gene expression. We also aimed to ensure this change in expression was generally greater than that of other SNVs for a given gene, to incentivize learning of the aSNV over other variant effects, and that the change in expression was the same for different genes, to simplify the pattern.

To support these goals, we first sought to have each aSNV, alone, explain a substantial fraction of the variance (R^2^) in gene expression across individuals for the gene it was paired with. We chose an arbitrarily large R^2^, 50%, to model the scenario where the aSNV is akin to an extremely strong causal eQTL SNV, whose genotype explains most of the differences in gene expression across individuals, and whose effect on gene expression is therefore expected to be greater than all or most other surrounding SNVs. The variance in gene expression (R^2^) explained by the aSNV genotype for a given gene depends on (1) the observed gene expression variance across individuals, (2) its MAF, and (3) its effect size. We therefore held the MAF of the aSNV constant (at 50%) for all genes, and standardized the variance in gene expression (detailed below) across individuals for each gene to hold (1) and (2) constant, and derived a constant effect size (3) for the aSNV (detailed below). Increasing the individual’s expression by a value of 2.0 enabled the aSNV to explain 50% of gene expression variance between individuals at a MAF of 50%, when introduced into personal genomes as a homozygous allele. We did not consider heterozygous alleles in this study, but the effect size of one copy of the effect allele is therefore 1.0.

To standardize each gene’s variance in gene expression across individuals, we divided the gene expression matrix by each gene’s standard deviation (across individuals), calculated separately between training, validation, and test donor splits to prevent leakage. This sets the standard deviation in gene expression to 1 for each gene (within each donor split).

While we set the MAF and R^2^ to be 50% to enable its outsized importance relative to other natural SNVs nearby, these were somewhat arbitrary choices. Therefore, we attempted modeling other MAFs and effect sizes (and therefore other R^2^ values), finding similar elastic net underperformance on the cross-individual expression task (**Supp. Fig. 8**)

We derived the aSNV effect size using the following formula, which assumes the effect allele will be introduced into personal genomes as homozygous (see above). In most setups, the R^2^ is set to 0.5 and the cross-individual variance in expression is standardized to 1. The variance in genotype across individuals resolves to 0.25 with a minor allele frequency of 0.5 and when the two genotype options are homozygous for the effect allele or homozygous for the other allele, yielding an effect size of 2.

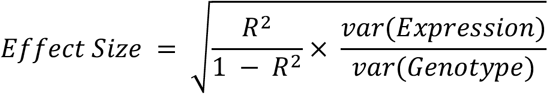

### Selection of Motifs and Motif Positions to Perturb with aSNVs

We started with JASPAR 2024 genome-wide motif scanning hits, obtained from the UCSC genome browser^29^ (JASPAR2024.bb from https://genome.ucsc.edu/cgi-bin/hgTrackUi?db=hg38&g=jaspar). We kept hits from JASPAR CORE motifs whose experimental data used to create the PWM derived from Homo sapiens and were within the 49,152bp TSS-centered windows we used to build personal genome sequences. We used the “score” field from this file to filter hits to a minimum score of 400 (p-value ≤ 10-4), to remove hits with weaker matches to the PWM. We refer to these as “candidate” motifs for receiving the aSNV.

From this pool, we chose specific motifs and relative motif positions to perturb based on experimental goals. For initial experiments using STAT1 (**Figs. 2a-c**), we chose one random STAT1 motif to perturb per gene and placed the aSNV in the 6^th^ position of the motif. These same motifs were perturbed in **Fig. 5a**. We also placed the aSNV in all other relative motif positions in **Supp. Fig. 11a**. In **Supp. Fig. 9a-b**, we overlapped ENCODE4 cCREs (encodeCcreRegistry.bb from the UCSC genome browser) with STAT1 motif hit coordinates. (minimum 1bp overlap), and picked one random site per gene, placing the aSNV at position 6.

In other experiments, we chose specific STAT1 motifs to perturb for particular experimental aims. In **Fig. 3a**, we chose STAT1 motifs to perturb that were either +/- 6kb from the gene’s TSS, or within 6kb from the edge of the same gene’s 49,152bp sequence window to investigate the effect of distance on learning variant effects. Among the motif candidates in each distance group for each train, validation, and test gene, we picked one random motif. We used the same train, validation, and test genes for both groups, such that the TSS distance of the perturbed motif was the primary difference.

In **Fig. 3b**, we chose STAT1 motifs to perturb that either came from regions with high or low pre-trained Enformer gradient attributions, as a proxy for the strength of pre-existing E2G links between the target gene and regions containing these motifs. We computed gradients of the gene expression prediction with respect to the hg38 reference input sequence, similar to the procedure we implemented in past work^16^. Briefly, we averaged Enformer outputs over the 3 TSS-overlapping bins, using the ‘CAGE:blood, adult, pool1’ (track 4950) track to get gene expression predictions. We next computed the 49,152bp x 4 input gradient matrix and retained the maximum absolute value along the nucleotide alphabet axis, yielding a 49,152bp x 1 matrix where each value represents the maximum absolute attribution of a nucleotide at that position. We then z-scored this matrix along the length axis, so each value represents the importance relative to the average among all positions. We defined regions with strong gradients as those whose gradient is at least 3 standard deviations greater than the mean for a given gene, while regions with weak gradients have a gradient value that is at most equal to the mean (i.e, z-score of 0). We required STAT1 motifs from both strong and weak gradient regions to be at least 6kb from the gene’s TSS, to focus on non-promoter motifs for which the model must form internal E2G links over distance. To control for distance and leave gradient attribution around the motif as the primary experimental variable, we attempted to match low gradient and high gradient motifs by their distance from the same gene’s TSS, while targeting the selection of 100 train, 50 validation, and 50 test genes with distance-matched strong and weak gradient motifs for model training and evaluation. Where possible, we matched their distance from the TSS to be within the same +/- 500bp window, but we matched strong gradient motifs with closest weak gradient motifs where this was not possible. We found 100 train, 47 validation, and 50 test genes with STAT1 motifs that fit these parameters. Over all 197 genes, the distance of strong and weak gradient motifs from the same gene’s TSS differed by a median of 2045bp within a 49kb sequence window.

In **Figs. 4a-b**, **5b** we trained 45 models, each by using aSNVs to perturb motif instances for one of 45 TFs that are expressed in Whole Blood. For each TF, we picked one random motif instance to perturb around each gene in this experiment’s train, validation, and test sets. We placed the aSNV in the highest information content position – where there is the strongest preference to one nucleotide above the others in the PWM – of the motif. We used the same train, validation, and test set genes to train each model.

### Selection of genes for training, validation, and test sets

In **Figs. 2a-c**, **5a** we started with a set of 300 genes that were used to fine-tune Enformer with personal genomes in our previous study^16^. Of these, 279 had a STAT1 motif within the 49kb window. We trained on these 279 genes. For 100 randomly selected genes, we set the aSNV MAF to 0% during training to model the learning of rare variant effects (**Supp. Fig. 4**). This left us with 279 training genes, and 179 aSNVs for training in our initial experiments. We then increased this to a dataset that is approximately 10x greater by identifying 2,725 genes, including the original 279 that are expressed in Whole Blood, that have a STAT1 motif within the 49kb window, and were in Enformer’s original train set. Finally, we included all possible genes that meet these criteria, yielding 12,111 train genes. For each of these three datasets, the same 100 genes are used to model rare variant effects, leading to 179, 2,625, and 12,011 training aSNVs. We started with 100 validation and 100 test genes from our previous manuscript, and used 92 and 88, respectively, that had STAT1 motifs within the 49kb window.

In **Fig. 3a**, we started with the 279 genes from above and identified those with both a STAT1 motif within 6kb of the TSS and 6kb from either the upstream or downstream window edge. This left us with 277 and 2713 train, 92 validation and 87 test set genes. To enable future rare variant analyses (as in **Supp Fig. 4**), we hid the aSNV for 100 of these genes, leaving us with 177 and 2613 training aSNVs. We did not train a model on the maximum possible number of genes (equivalent to the 12,011 aSNV model) to conserve computational resources. In **Fig. 3b** we used 100 train genes, 50 validation, and 50 test set genes, chosen based on selection criteria for perturbed STAT1 motifs within their window (see **Selection of Motifs and Motif Positions to Perturb with aSNVs**).

In **Figs. 4a-b**, **5b** we trained each of 45 models (one per TF motif) on the same 300 train genes, 100 validation genes, and 1,183 test set genes. We picked a random set of 300 train and 100 validation set genes for which at least one instance of each of these motifs was present within the 49kb sequence window. We selected every gene in Enformer’s original test set that was expressed in Whole Blood and where at least one instance of each of these 45 motifs was present in the 49kb window to obtain our test set of 1,183 genes.

### Analysis of sequence context variability through each motif’s information content and palindrome score

To describe the aSNVs sequence context variability when placed in different TF’s motif instances, we extracted a 30bp window from the hg38 reference centered on the motif instance around every gene in the train, validation, and test sets that was selected to receive the aSNV. Instances on the (-) strand were reverse complemented. We computed a position frequency matrix (PFM) over these instances by averaging their one-hot encodings over these 30bp windows. We repeated this or each of 45 motifs we used to train models in **Figs. 4a-b** and **5b**.

We calculated positional information content from this PFM by taking 2 - H, where H is the Shannon entropy of the nucleotide distribution at that position. This was done over all 30 positions in the PFM, including those that flank the core motif, and the mean over the 30 positions was taken. Because the window extends past the core motif, this value reflects the information content of the core motif, its length, and the consistency of the flanking nucleotides (typically random, low information content). Longer, higher-information content motifs, which reduce sequence context variability the most, are upweighted in this way.

To compute each motif’s palindrome score, we used only the portion of the PFM covering the motif itself, excluding the flanking sequence that makes up the rest of the 30-bp window. The palindrome score summarizes how different the motif looks from its own reverse complement. We subtracted the reverse-complement PFM from the original PFM. At each position, this gives the difference in frequency of each nucleotide between the two orientations. If a position matches its counterpart in the reverse complement, all four differences are zero. Otherwise, some nucleotides are more frequent in one orientation than the other, and we recorded the largest such difference as a negative value, ranging from 0 (identical) to −1 (no nucleotides in common). Values in between indicate partial mismatches: for example, −0.3 means that one nucleotide occurs in 30% more of the motif instances in the reverse-complement orientation than in the original. Summing these values across all positions of the motif gave the palindrome score. A score of 0 indicates a perfect palindrome, in which the dominant nucleotide in the motif’s forward orientation is identical to that of its reverse complement, and increasingly negative scores indicate greater differences.

For each of the 45 TFs, we recorded the mean cross-individual correlation across the 1,183 test set genes (HOGP) achieved by the Enformer model fine-tuned on its aSNV dataset. We regressed this performance on the mean information content and palindrome score of each motif using the statsmodels^30^ ordinary least squares implementation, after z-scoring both features. In **Fig. 5b**, we report this mean cross-individual correlation for each TF on the y-axis, and the linear model’s fitted values using these two features on the x-axis.

### Learning rare aSNVs

To model the scenario where rare variants are not observed during training but appear in held-out individuals, in genes that were observed during training, we set the MAF of the aSNV to 0% for a subset of 100 genes in **Figs. 2a-c** and **Supp. Fig. 4**. We then set the MAF back to 50% for individuals in the validation set, and we investigated whether there is a difference in Enformer’s ability to learn unseen variants around genes observed during post-training, versus unseen variants in totally held-out loci. We then compared cross-individual correlation, among HOP in the validation set, to see whether learning of the two plateaus at different points during training. For this reason, in **Figs. 2a-c**, we train on 100 additional genes than aSNVs (e.g., 179 aSNVs seen around 279 genes).

### Enformer architecture when fine-tuning on aSNVs

We fine-tuned Enformer the pytorch implementation of Enformer (https://github.com/lucidrains/enformer-pytorch) using a similar architectural and training hyperparameters as our previous manuscript^16^. Briefly, we updated all weights, used 49kb sequences, and replaced the human and mouse output heads with a linear layer. We kept one of the output bins overlapping the TSS (bin 192 out of 384 for a 49kb DNA sequence) and computed the loss (mean-squared error between predicted and observed individual expression) against this value during training. We used the AdamW optimizer, 5e-6 learning rate, bf-16 mixed precision, and a batch size of 28. For most experiments, we trained using 4 H100 GPUs, using pytorch-lightning’s (https://lightning.ai/docs/pytorch/stable/) implementation of the DataDistributedParallel strategy. When doing this, we multiplied the learning rate by the number of GPUs being used to account for the increase in effective batch size. Each example in a batch corresponds to a different individual-gene pair. After each epoch, we iterated through genes and individuals in the validation set. We monitored the per-gene cross-individual correlation, averaged over validation set genes, and stopped training if this value did not improve after 20 epochs. We kept the best checkpoint on this metric.

### Choice of individuals for training and evaluation

Starting with 670 individuals with Whole Blood RNA-seq and WGS data available from GTEx v8, we applied the same 80%/10%/10% train, validation, and test set splits as our previous manuscript^16^, yielding to 532, 67, and 71 individuals in each set. The same splits are used to train every model; where model replicates were trained, they differ based on random seed.

### Predicting aSNV effect sizes

We implemented *in silico* mutagenesis to calculate aSNV effect sizes. Briefly, for an aSNV around a given gene, we perform two forward passes using that gene’s TSS-centered, 49kb reference genome sequence (hg38). At the aSNV position, we modified one of these sequences to contain the effect allele and one to contain the other allele, and took the difference in predictions between the two as the effect size. We repeated this with the reverse complement of these two sequences, and averaged the effect size over the two orientations.

### Fitting elastic net baselines

We fit elastic net models using ElasticNetCV from scikit-learn^31^ as we have done previously^16^. Briefly, we trained one model for each gene, using a max_iter of 2000. We matched train and test individuals with our fine-tuned Enformer models. We fit these elastic nets using a matrix of genotypes (0, 1, or 2) for each SNV observed in the same individuals used to fine-tune Enformer models for a given gene. Thus, the genetic variation elastic nets and Enformer see during training and evaluation are identical. Gene expression values for a given individual and gene were the same as used for Enformer fine-tuning and evaluation. To match the aSNV setup, the elastic net genotype matrices also included the aSNV genotype for each individual, matching what Enformer saw. We fit and evaluated these elastic nets two ways: (1) using all observed SNVs and the aSNV or (2) the aSNV only. Unless otherwise stated in the text, elastic net baselines come from (2). In **Fig. 2a**, the elastic net baseline used for the pre-trained Enformer model that was evaluated on personal genomes without aSNVs was also not fit or evaluated using aSNVs.

### Calculation of cross-individual correlation

We obtained a vector of true gene expression values from 71 individuals in the test set. These gene expression values include aSNV-induced changes to gene expression, if the individual had the “effect” allele. We then obtained a vector of predicted gene expression values by passing in their personal genome sequence for fine-tuned Enformer. We repeated this with both forward and reverse complement orientation sequences, averaging predictions for each individual and gene over the two. For the elastic net baseline, we passed in either their aSNV genotypes, or a matrix of their aSNV and observed SNV genotypes (see **Fitting elastic net baselines**), depending on the type of baseline used. We used these vectors of observed and predicted expression and took the Pearson correlation between the two to obtain the cross-individual correlation for each model type. When multiple fine-tuned Enformer replicates were trained (different random seeds), we averaged cross-individual correlations, as described in figure captions.

### Analysis of fine-tuned Enformer’s CNN and Transformer embeddings

After fine-tuning Enformer on an aSNV dataset, we load the trained model from its checkpoint. We then passed in that model’s train and test sets and retrieved embeddings from the final CNN and final transformer layers. These datasets appeared identically as during training and evaluation of the model. In particular, aSNV positioning, expression values, and aSNV genotypes were unchanged. From these two embeddings, we extracted the bins along the sequence axis overlapping the aSNV and the TSS, as well as a bin on either side of them. For the four embedding types (2 layers x 2 positions) that left us with a 3×1536 matrix for each (3 bins, 1536 channels). We averaged over the 3 bins, yielding one 1536-dimension vector per gene and individual.

We then fit a separate elastic net on each of the four embedding types, using 1536-dimension embedding vectors from each gene and individual in the train set to predict that individual’s gene expression value. This gene expression value is identical to what Enformer saw for this gene and individual during fine-tuning and reflects the aSNV genotype. The fitted elastic net is then used to form predictions using 1536-dimension embedding vectors from the test set (fine-tuned Enformer’s HOGP) as input. The resulting gene expression predictions were used to form cross-individual correlations for each gene, and these values were compared against separate elastic net baselines that take genotype as input and were used to benchmark HOGP accuracy throughout this manuscript (see **Fitting elastic net baselines**).

Regression models in **Figs. 4a-b** were fit and evaluated separately on the same 45 fine-tuned Enformer models reported in **Fig. 5b**.

### Fine-tuning Enformer with aSNVs that perturb modified STAT1 motifs

We first generated a random 1000bp left-flank and a random 1000bp right-flank to place around naturally occurring STAT1 motifs and trimmed each to 3,6, or 9bp flanks, giving STAT1 motifs of 15, 21, or 27bp (after accounting for the original 9bp core). We ensured the positioning of nucleotides in these trimmed flanks remained consistent (e.g., the 3bp flank is the inner part of the 9bp flank and remains closest to the motif), such that the only difference between the three groups were the added nucleotides. We started with 1000bp flanks to enable testing of additions longer than 9bp, if desired, while maintaining this consistency.

For every STAT1 motif that was a candidate for receiving the aSNV (i.e, those with at least a score of 400, see **Selection of Motifs and Motif Positions to Perturb with aSNVs**) we replaced the reference bases around it with these left and right flanks, including candidate STAT1 motifs that did not receive the aSNV. The same left flank sequence was used at every motif instance, and the same right flank sequence was used at every motif instance. For STAT1 motifs present on the (-) strand, the flanks were reverse complemented and left-right swapped to maintain orientation. After introducing the aSNV, we saved these sequences to HDF5 files that were then loaded during training and evaluation, as performing all of these modifications to the sequence on the fly led to a CPU bottleneck. Introducing these added flanks were meant to test whether decreasing sequence context variability by increasing motif length and information content led to more data efficient learning of aSNV effects. We extended every candidate STAT1 motif, not only the one with the aSNV to ensure the model could not find the aSNV by recognizing the flank alone, rather than distinguishing one aSNV-perturbed motif from many unperturbed ones that are otherwise identical.

Besides this modification to perturbed and unperturbed candidate STAT1 motifs, the rest of the experiment was identical to that of the 179 aSNV training experiment in **Fig. 2**: the same STAT1 motifs were perturbed with the same aSNVs in the same positions, leaving the STAT1 modification as the primary difference.

### Data augmentation by sliding aSNV-perturbed STAT1 motifs around the TSS

For each training example, we took the aSNV-perturbed STAT1 motif, as well as the 50bp of flanking sequence on each side and moved it by a random distance to another location on the same side of the TSS, requiring that it be at least 1kb from the TSS. As the perturbed STAT1 motif slides from its original position to the new one, the nucleotides it slides past are displaced 109bp (50bp flank + 9bp STAT1 motif + 50bp flank) in the opposite direction, landing near their original position and with the same neighboring sequence. A new STAT1 location was drawn independently during every training example, such that the perturbed STAT1 motif appears in many different locations around the target gene during training. We slid aSNV-perturbed STAT1 motifs only during training, and not during validation or testing. Besides the sliding of STAT1 motifs, the rest of the experiment was identical to that of the 179 aSNV and 2,625 aSNV training experiments in **Fig. 2**, leaving the data augmentation strategy as the primary difference

### Counting fm-eQTLs that overlap ChIP-seq peaks and motifs

We obtained JASPAR 2024 motif sites (see **Selection of Motifs and Motif Positions to Perturb with aSNVs)** and ENCODE4 TF rPeak clusters (TFrPeakClusters.bb) from the UCSC genome browser. We again restricted the analysis to JASPAR CORE TFs whose experimental data used to create the JASPAR PWM came from Homo sapiens. We filtered these rPeak clusters to include those whose experiment list included ChIP-seq from GM12878 or K562 cells (blood-relevant cell lines). For each TF, we intersected its JASPAR motifs with its own rPeaks, keeping instances where the JASPAR motif overlaps the same TF’s rPeak by at least 1bp. For added leniency, we included all JASPAR annotated motifs for each TF, including those that most weakly match the PWM, as well as all rPeak clusters regardless of signal strength. We then intersected these overlaps with Whole Blood fm-eQTLs (CAVIAR^32^ posterior inclusion probability > 0.9) from GTEx, requiring that the fm-eQTL fall within the JASPAR motif that is within the larger rPeak. This left us with 137 TFs. For each TF, we then counted the number of fm-eQTLs that fell within any of its rPeak-overlapping motifs.

### Fine-tuning Enformer on rare multi-tissue expression outlier SNVs

From GTEx v8^21,33^, we obtained median cross-tissue z-scores (per person and per gene; gtexV8.eOutlier.stats.globalOutliers.removed.txt.gz). We attempted to use these z-scores to fine-tune Enformer on rare SNVs with large effects on expression that persist over multiple tissues, similar to the promoterAI^15^ approach. As elsewhere in this study, we used 49,152 bp sequence windows centered on each gene’s TSS, and paired each SNV with every gene whose window contained it. For each SNV and gene pair, we performed a t-test comparing the distributions median z-scores among carriers and non-carriers. We selected SNVs with a nominal *p* < 0.01 from this t-test for training, and used the average z-score among carriers as the estimate of the SNVs directional effect on expression. We filtered these SNVs to only include those outside of coding sequences. Promoter “Proximal” SNVs were defined as those within 500bp upstream or downstream of a gene’s TSS, while “Distal” SNVs were defined as those at least 1kb away. Splitting SNVs into train/validation/test splits based on Enformer’s original splits, we had 5,447/388/586 train/validation/test “Proximal” SNVs and 122,293/8,601/13,764 “Distal” SNVs. The “Proximal + Distal” set was the sum of the two.

We trained a ‘TwinNet’ Enformer architecture, which others observed to work effectively for similar tasks^15^. Specifically, for each SNV, we perform an Enformer forward pass with both a reference (hg38) genome sequence, and a sequence that contains the alternate allele but is otherwise identical. We retrieve the embeddings from both following Enformer’s trunk. We then subtract the reference embedding matrix from the alternate allele embedding matrix, and average the result over the sequence and channel axes. Finally, we pass the resulting scalar through a tanh activation function. We froze all weights except the final layer of Enformer’s trunk. We used an AdamW optimizer, bf-16 mixed precision, and 4 H100 GPUs, managed by pytorch-lightning’s implementation of the data distributed parallel strategy. Each GPU received a batch of 14 SNVs, and the learning rate was 2e-3. Our loss function was the mean squared error between predicted and observed effect sizes for each SNV-gene pair.

After each training epoch, we evaluated the model on unseen outlier SNVs found around genes in the validation genes. Validation genes included any gene from Enformer’s original validation set that included at least one outlier SNV. We monitored the loss over this validation set, and stopped training if the loss did not improve after 20 epochs, and the best checkpoint was kept.

We evaluated resulting models on various promoter variant-effect benchmarks from the promoterAI manuscript, by correlating the predicted effects of SNVs in each benchmark to the true effect size estimates we obtained from the authors. We omitted SNVs from these benchmarks if their ‘consequence’ value was ‘none’. We removed SNVs in each benchmark if they were in the train or validation set of any of the TwinNet Enformer models. We also evaluated these models on promoter-distal (>1kb from TSS) multi-tissue fm-eQTLs, using those with posterior inclusion probability (PIP) > 0.9 in at least two tissues. We set the true effect size value of these multi-tissue fm-eQTL to the mean slope over the tissues with PIP > 0.9. TwinNet Enformer predictions are averaged over forward and reverse complement sequence orientations. We also evaluated Enformer on the same SNVs. To compute a predicted effect size, we performed a forward pass with the reference and alternate allele sequences, obtaining a 384 x 5,313 matrix from the Enformer human head. We subtracted the reference matrix from the alternate allele matrix, and averaged over the sequence and channel axes. As others observed^15^, we found averaging over the entire sequence and all channels optimized performance on these variant effect benchmarks.

To evaluate TwinNet Enformer and pre-trained Enformer on personal genome sequences, we used 100 test set genes from our previous manuscript and compared their predictions to individuals’ gene expression values in GTEx Whole Blood. We picked these genes because an elastic net achieves a cross-individual correlation of at least 30%. As the TwinNet requires two sequences as input, we formed these predictions by passing in an individual’s personal genome sequence paired with the reference genome sequence from the same locus. For Enformer, we averaged outputs over the 3 TSS-overlapping bins, using the ‘CAGE:blood, adult, pool1’ (track 4950) track, finding this approach led to slightly improved predictions relative to the all-channel, all-bin aggregation employed for the promoterAI benchmarks.

## Acknowledgements

We thank Nilah Ioannidis, Tony Capra, Sandy Floren, Ruchir Rastogi, Anshul Kundaje, Yun Song, and David Kelley for discussion and feedback. We thank Gherman Novakovsky for code and advice regarding fine-tuning TwinNet Enformer on multi-tissue outlier variants, as well as for benchmarking data. Lastly, we thank Peter Koo for helpful feedback on the writing of this manuscript, as well as advice regarding data augmentation strategies.

## Declarations

### Code availability

Code to implement LAMP can be found at https://github.com/shirondru/LAMP.

