## Supplemental Figures for "LAMP sheds light on how sequence to function models learn variant effects during post-training"

### Supplementary Figures

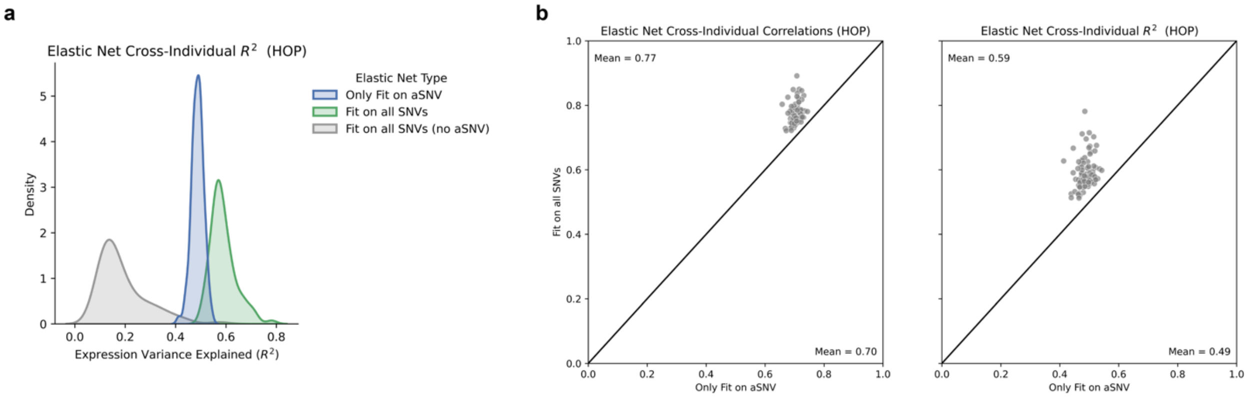


**Supplementary Figure 1: The aSNV's outsized expression effect dominates natural SNV contributions and explains most of the total genetically-regulated component of gene expression**

**A)** $R^2$ values achieved by three elastic net models, fit on genes from fine-tuned Enformer's test set and evaluated on held-out people. One model predicts each individual's gene expression using only each individual's aSNV genotype (blue), another also considers naturally observed SNVs in the same sequence window around each gene (green), and the last one considers only natural SNVs. The first two are fit and evaluated using expression values modified to reflect to the aSNV effect size, the last one is fit and evaluated on unmodified values. The expression variance explained by the aSNV is much greater than that of all of the natural SNVs in the same loci. **B)** Cross-individual correlation (left) and $R^2$ (right) values achieved by two different elastic net models, fit on genes from fine-tuned Enformer's test set and evaluated on held-out people. One model predicts each individual's gene expression using only each individual's aSNV genotype (X-axis), the other also considers naturally observed SNVs in the same sequence window around each gene (Y-axis). The latter forms slightly more accurate gene expression predictions because it also captures the importance of natural SNVs, but the small overall difference reflects the outsized importance of aSNVs to individual gene expression predictions, and that its genotype alone accounts for nearly all of the cross-individual variation in expression these models can explain. As fine-tuned Enformer underperforms the linear baseline formed by using just aSNVs (X-axis, **Fig. 2a**), this underperformance cannot be explained by a failure to learn natural variant effects, and suggests it is instead caused by incomplete fine-mapping of aSNV effects (consistent with our past findings[10]).

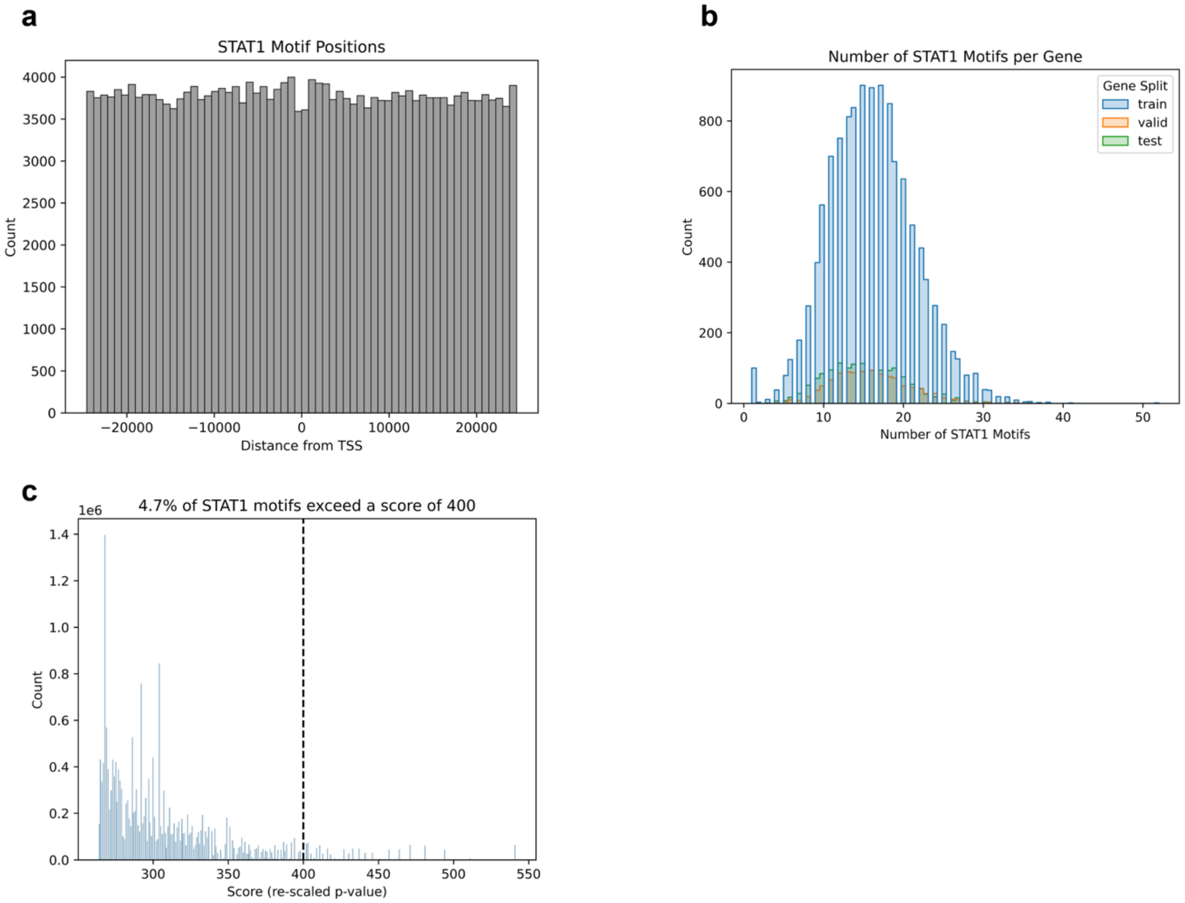


**Supplementary Figure 2**: **STAT1 Motif Properties**
**A)** The relative position of strongly matching candidate STAT1 motifs (**Methods**) relative to the TSS of 14,696 genes that are expressed in Whole Blood and within 49kb of a nearby protein-coding gene. **B**) The number of STAT1 motifs within the 49kb window around each of these genes, separated by whether they fall in the train, validation, or test set using Enformer's original splits. We separate genes using the same splits in this manuscript. The median gene has 16 STAT1 motifs within a 49kb window around it. **C**) Distribution of all JASPAR-annotated STAT1 motif scores, with a vertical line at 400 (corresponding to $p = 10^{-4}$**).** Thresholding at 400 leaves behind 4.7% of the strongest matching motifs, used elsewhere throughout this manuscript.

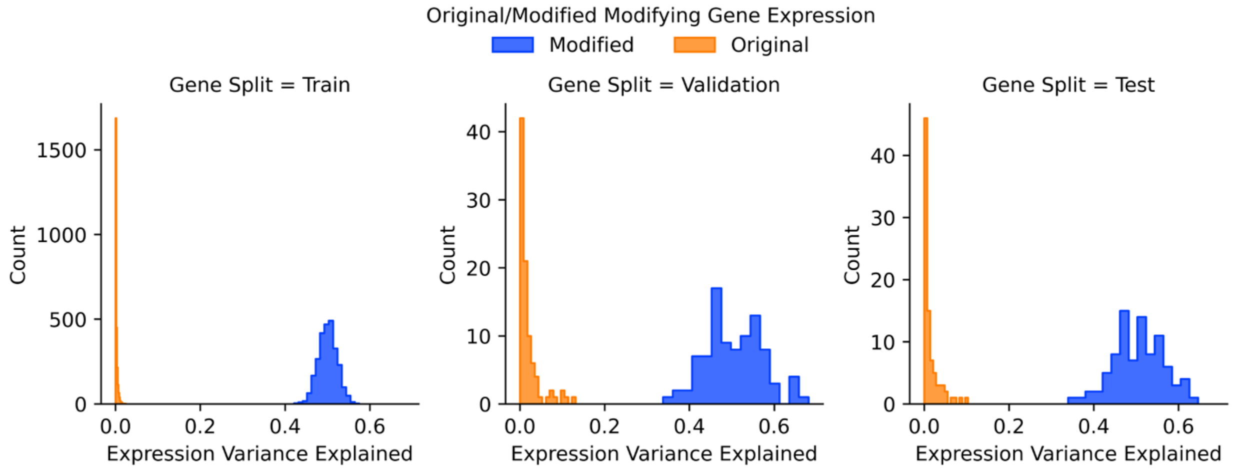


**Supplementary Figure 3: The aSNV explains 50% of expression variance**
For genes in the train, validation, and test sets, we regressed expression on aSNV genotype across individuals from the train, validation, and test sets, using either original GTEx Whole Blood expression (orange) or their expression after introducing the aSNV effect (blue). This regression was done one gene at a time, and we plotted the variance explained by the aSNV along the x-axis. As aSNV genotypes are assigned to individuals randomly (at 50% MAF), they explain close to none of the variance in the original expression, but close to the target of 50% after introducing the aSNV effect. Variance explained varies more for validation and test genes because fewer individuals are included in these sets (532 train individuals, 67 validation individuals, 71 test individuals).

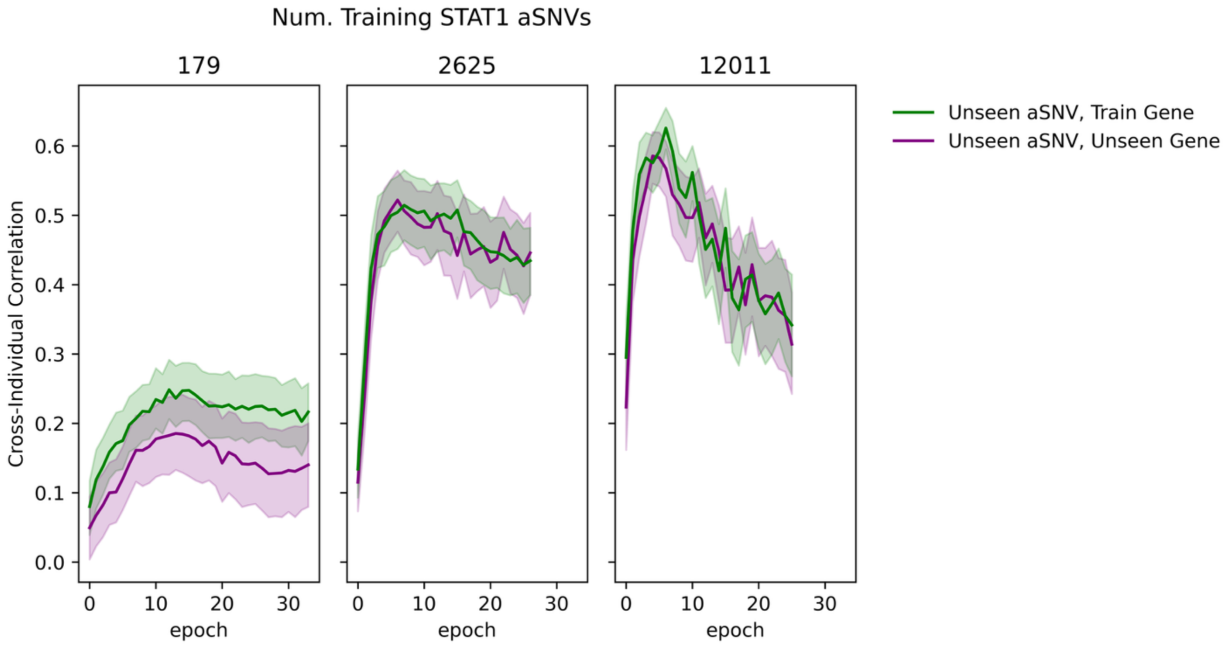


**Supplementary Figure 4: Simulating the learning of rare variant effects with LAMP**
Cross-individual correlation among HOP in the validation set and train genes (green) or validation genes (purple) versus training epoch, for models being fine-tuned on different dataset sizes. The aSNV was held-out of these train genes (green) during training, but reintroduced in these individuals in the validation set. These results suggest unseen variants in loci observed during training are learned at approximately the same rate as unseen variants in totally held-out loci. We speculate that the 179 training aSNV model achieved higher performance on train genes relative to unseen ones because this model is making use of natural variants also observed around these genes that can help explain gene expression differences in held-out people. This disappears in the models trained on more data, as they learn to upweight the aSNV more (as it explains most of the gene expression differences) and attend to other variants less.

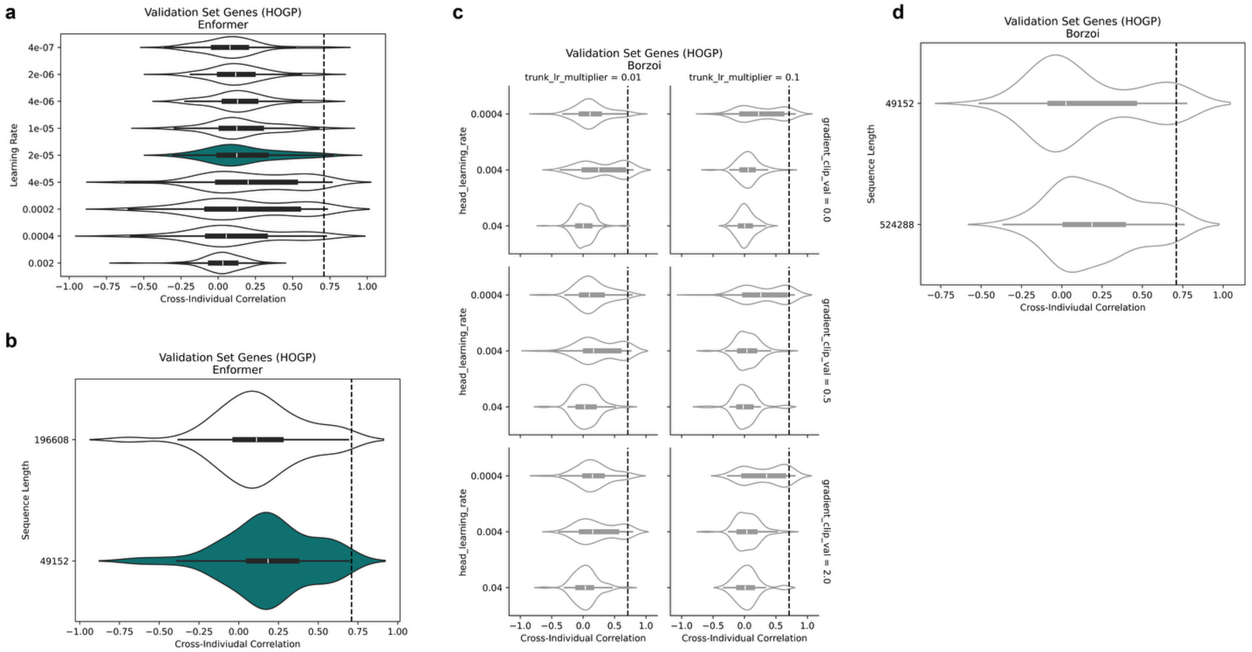


**Supplementary Figure 5: Elastic net underperformance holds when modifying hyperparameters and switching to Borzoi**

**A)** HOGP performance using genes and individuals from the validation set after varying learning rate during training. In teal is the learning rate used elsewhere in this manuscript. **B**) Validation set HOGP after training and evaluating on 196kb sequences or 49kb sequences. In teal is the sequence length used elsewhere in this manuscript. In **A-B**, we tested the robustness of the hyperparameters we previously tuned for Enformer in a prior manuscript, using validation set genes and individuals to avoid overfitting on the test set. None of the tested values rescued elastic net underperformance relative to those used elsewhere in this manuscript, which are the ones we previously found to be optimal[16]. **C**) Validation set HOGP performance using a pytorch borzoi implementation from Flashzoi[34] repository. We used the original version of Borzoi (replicate 0) without Flash Attention, starting from its pre-trained weights, using 49kb sequences. We replaced the Borzoi heads with a linear layer to map the embeddings to a single output track. We varied the learning rate of this output head (“head_learning_rate”). We also varied the learning rate of the trunk by setting it to be a fraction of the head_learning_rate, because the trunk we pre-trained and the head was randomly initialized. We also tested clipping the gradient’s global norm to different values. All combinations underperform the elastic net by a wide margin, as with Enformer. **D)** Same as (**B**) but with Borzoi, using 49kb and 524kb sequences. All Enformer models here are trained on 179 aSNVs and evaluated on 92 genes from the validation set (each with one aSNV). All Borzoi models are trained on 178 aSNVs and evaluated on 91 genes, with one dropped from each set because these genes sit near a chromosome boundary edge, and generating 524kb TSS-centered sequences for them was impossible. Dashed vertical lines represent elastic net baselines.

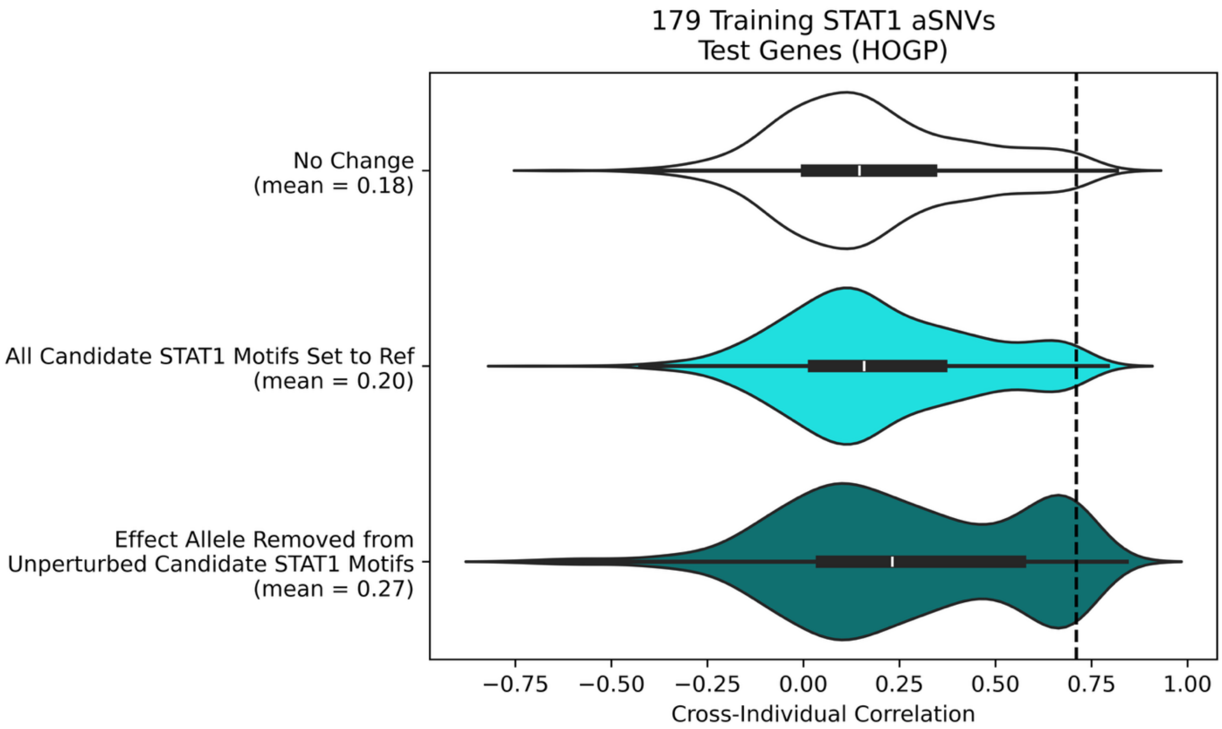


**Supplementary Figure 6: Controlling for LAMP confounders**
Cross-individual correlation on HOGP after making LAMP modifications to account for possible confounders. All models were trained on 179 STAT1 aSNVs, and the “No Change” group is the same as the 179 aSNV group in **Figs. 2a-b**. We accounted for the possibility of genetic variation in STAT1 motifs (including those with the aSNV and those without it) by taking the trained 179 STAT1 aSNV model from **Figs. 2a-b** and re-evaluating it on HOGP after removing genetic variation from all candidate STAT1 motifs by replacing their sequence with that of the hg38 reference. We also accounted for the natural presence of the effect allele (G at position 6) in STAT1 motifs that did not receive the aSNV. This may confound the model during training, as these G nucleotides are not associated with a change in expression. We controlled for this by replacing these G nucleotides with the other allele (C) in candidate STAT1 motifs. We then re-trained and re-evaluated on HOGP. We found these are minor confounders that modestly hinder learning of aSNV effects, but correcting them does not rescue elastic net underperformance.

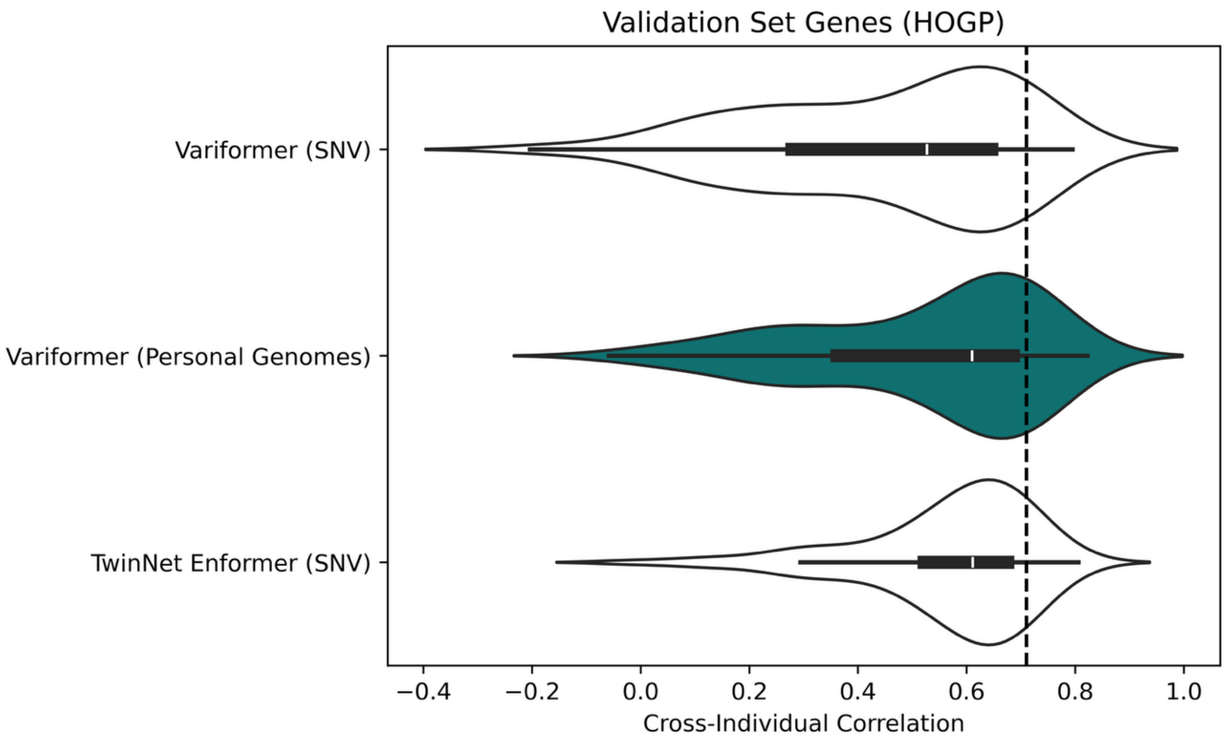


**Supplementary Figure 7: Elastic net underperformance holds when training directly on aSNV effects**

HOGP performance using genes and individuals from the validation set when training on personal genomes (as done elsewhere throughout this manuscript, teal) versus training on individual variants using the same Variformer[16] architecture or a TwinNet Enformer architecture, used in Jaganathan et al.[15] (**Methods**). For the Variformer variant model, each training example is the reference sequence carrying one of four alleles: the aSNV effect allele, the aSNV other allele, a natural SNV's alternate allele, or a natural SNV's reference allele (unmodified reference sequence). The corresponding target expression values are, respectively, the gene's mean expression plus the aSNV effect size, the gene's mean expression, the mean expression of carriers, and the mean expression of non-carriers. aSNV effect sizes are the same as those used when training on personal genomes. Natural background SNVs were randomly sampled from GTEx variants observed in training-set individuals, and included for training to prevent overfitting and memorization of the aSNV effect. For TwinNet models, the target is the difference between the two alleles' target values. All models use a mean-squared error loss, 49kb sequences, and are trained on the same 2625 aSNVs and evaluated the same 92 validation set genes (HOGP; one aSNV per gene). We train variant models on naturally occurring background SNVs because we found that, when not included, models will overfit to the aSNV effect size, and predict similarly large effects for any arbitrary variant.

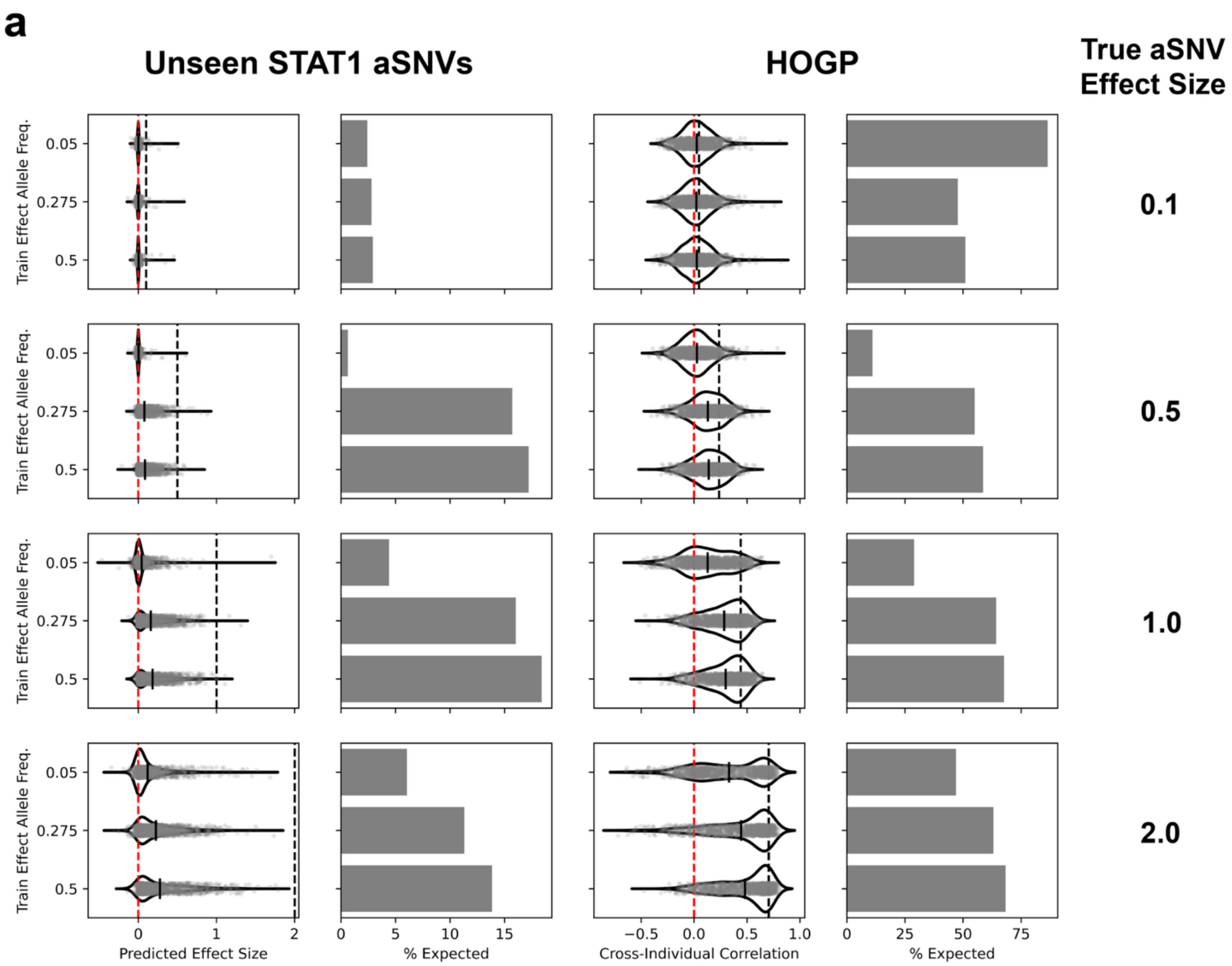


**Supplementary Figure 8**: **Modifying MAF and effect size of aSNV during training**
While the effect size and MAF of the STAT1 aSNV was set to 2.0 and 50% throughout the rest of this manuscript (**Methods**) we evaluated the outcome after varying these parameters. We attempted aSNV MAFs of 5%, 27.5% and 50%, and true effect sizes of 0.1, 0.5, 1.0, and 2.0. We set the MAF to 50% for individuals in the test set to ensure comparability within a given effect size, and to ensure low MAF groups were not disadvantaged during personal genome prediction evaluations. Left column shows predicted SNV effect sizes as the true effect size in these individuals is varied (panel rows) and as the MAF during training varies (rows within a panel). Dashed black line represents the true effect size Next column shows the mean predicted effect size by each model as a percentage of the true aSNV effect size. Next column shows HOGP predictions for the same test genes for which the aSNV effect size predictions were made, with vertical bars again denoting the mean and dashed black line representing the elastic net baseline performance (elastic nets fit and evaluated on datasets with the same MAF/effect size parameters). Last column shows the mean HOGP accuracy from each of these models as a percentage of the elastic net accuracy. Modifying effect sizes leads to similar underestimated effect sizes and underperforming HOGP predictions except for the lowest (effect size = 0.1) group, where evaluations were less reliable because the expected value was so close to 0.

Decreasing MAF during training led to substantially worse learning and generalization of the aSNV. All models were trained on 2,625 aSNVs.

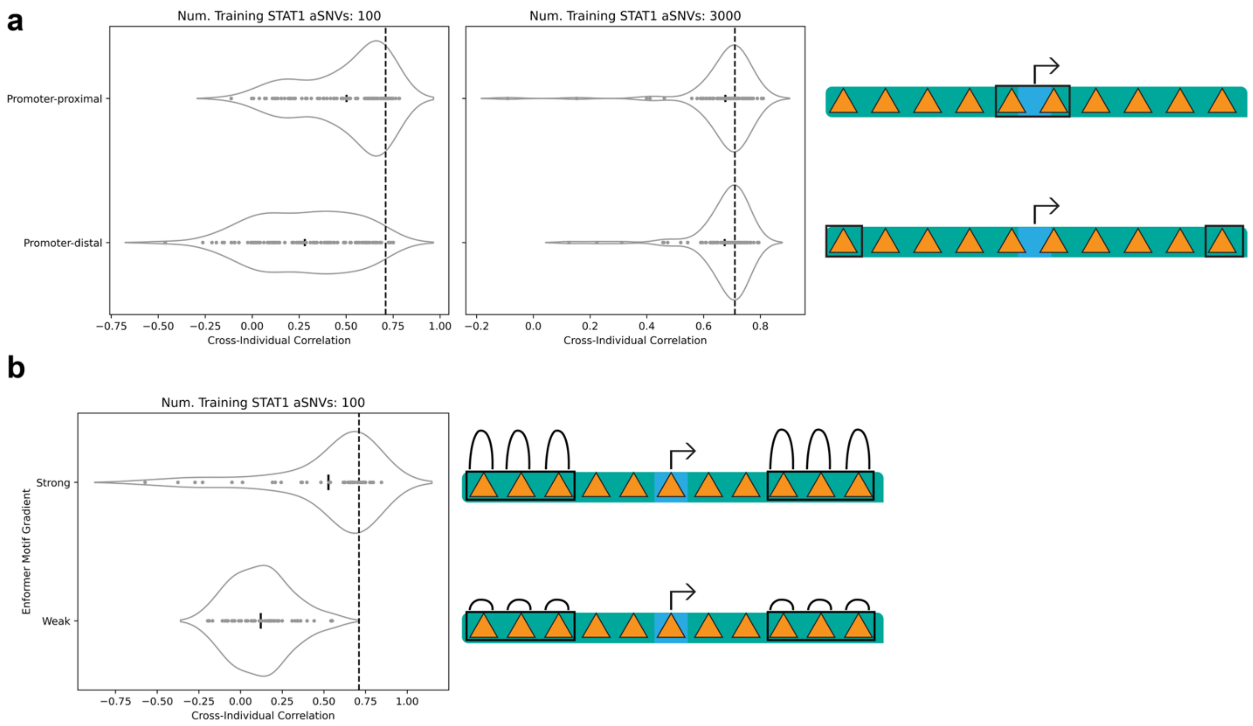


**Supplementary Figure 9: Enhancer-to-gene link issues persist when perturbing cCRE-overlapping STAT1 motifs with aSNVs**

(**A**) and (**B**) are similar to **Fig. 3a-b**, respectively, except the motifs perturbed with aSNVs here overlap cCREs.

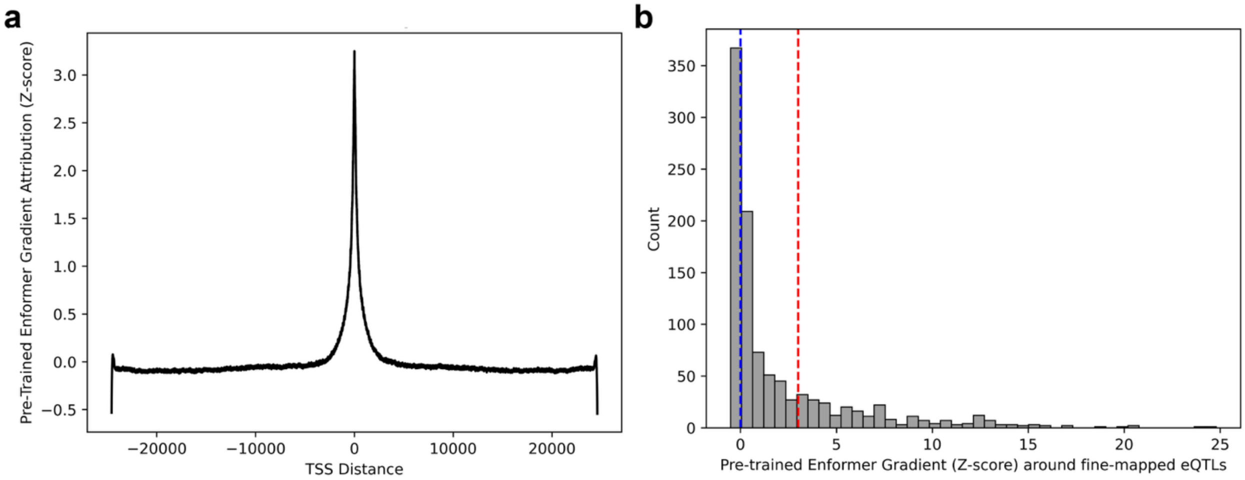


**Supplementary Figure 10: Enformer downweights TSS-distal regions, including those overlapping fm-eQTLs**

(**A**) Pre-trained Enformer's gradient attribution in the 49kb window around 14,696 genes expressed in Whole Blood. Gradient value is z-scored (**Methods**) to show the importance of each nucleotide along the sequence relative to all others, for each gene. Gradient attribution decreases sharply with distance from the TSS. (**B**) Gradient values (z-scored) around 1,023 Whole Blood fm-eQTLs SNVs found around 845 genes that overlap a JASPAR motif. Gradient values are the average gradient across the motif overlapping the fm-eQTL. Red and blue dashed lines represent the gradient z-score thresholds used to threshold motifs in strong and weak attribution regions, respectively, in (**Fig. 3b** and **Supp. Fig. 9b; Methods**). Most fm-eQTLs are in motifs with weak attributions.

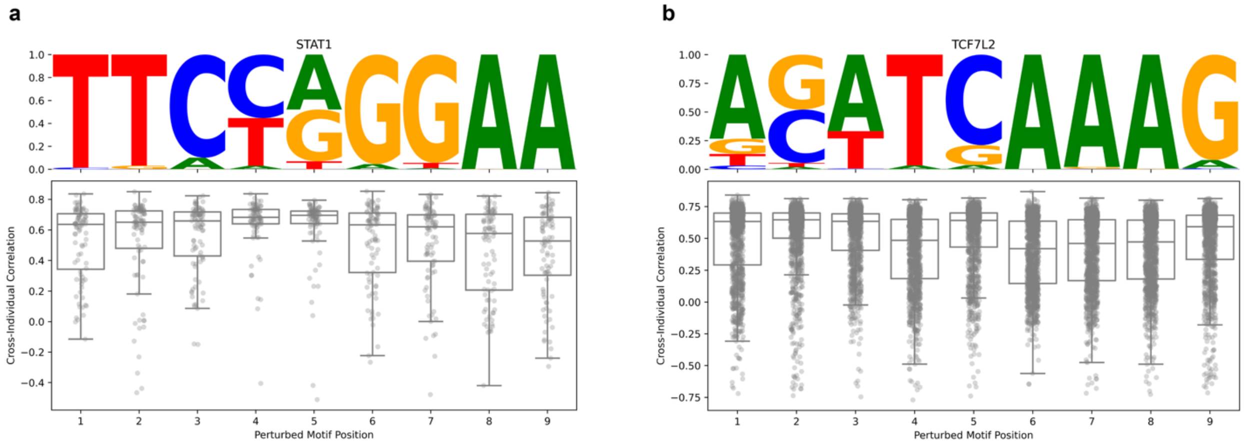


**Supplementary Figure 11: HOGP accuracy after perturbing different motif positions with aSNVs**

**A)** The STAT1 PWM (JASPAR motif MA0137.4) (top) and HOGP accuracy after fine-tuning Enformer while placing the STAT1 aSNV in each position, for a total of 9 models (bottom). Each point is a gene from the test set (HOGP). Results from **Fig. 5a-b** suggest learning of aSNVs is sensitive to sequence context variability. Consistent with this, HOGP performance increases when the aSNV is placed in lower IC positions, which decreases sequence context variability by removing a low IC position and replacing it with the aSNV. (**B**) is the same as **A**, but with TCF7L2 (JASPAR motif MA0523.2).

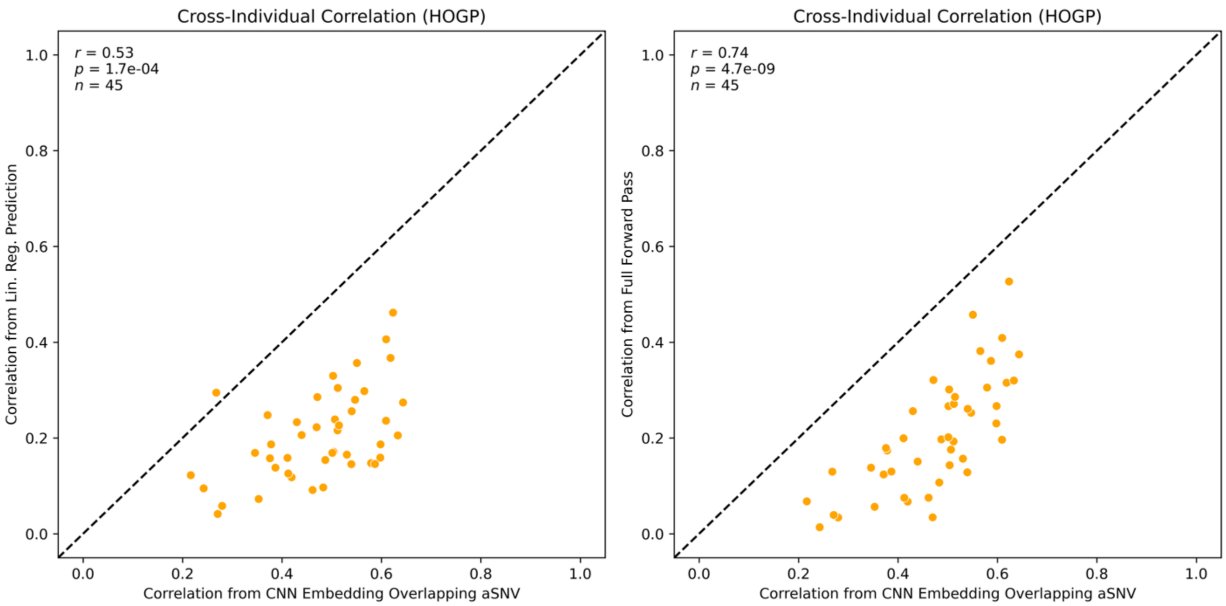


**Supplementary Figure 12: Sequence context variability helps explain CNN aSNV detection issues**

Cross-individual correlation (HOGP) after fitting regression models to predict individual gene expression from CNN embeddings overlapping the aSNV (from **Fig. 4**) is on the X-axis. Each point represents a different aSNV-perturbed motif during Enformer fine-tuning. Y axis is the cross-individual correlation from the same model's full forward pass (right, identical to orange points in **Fig. 4b**) or a linear regression model fit to predict these HOGP values based on each motif's IC and palindrome score (**Fig. 5b**). As we treat IC and palindrome score as a proxy for sequence context variability, this suggests sequence context variability helps explain the ability of the CNN module to distinguish aSNV genotypes after fine-tuning.

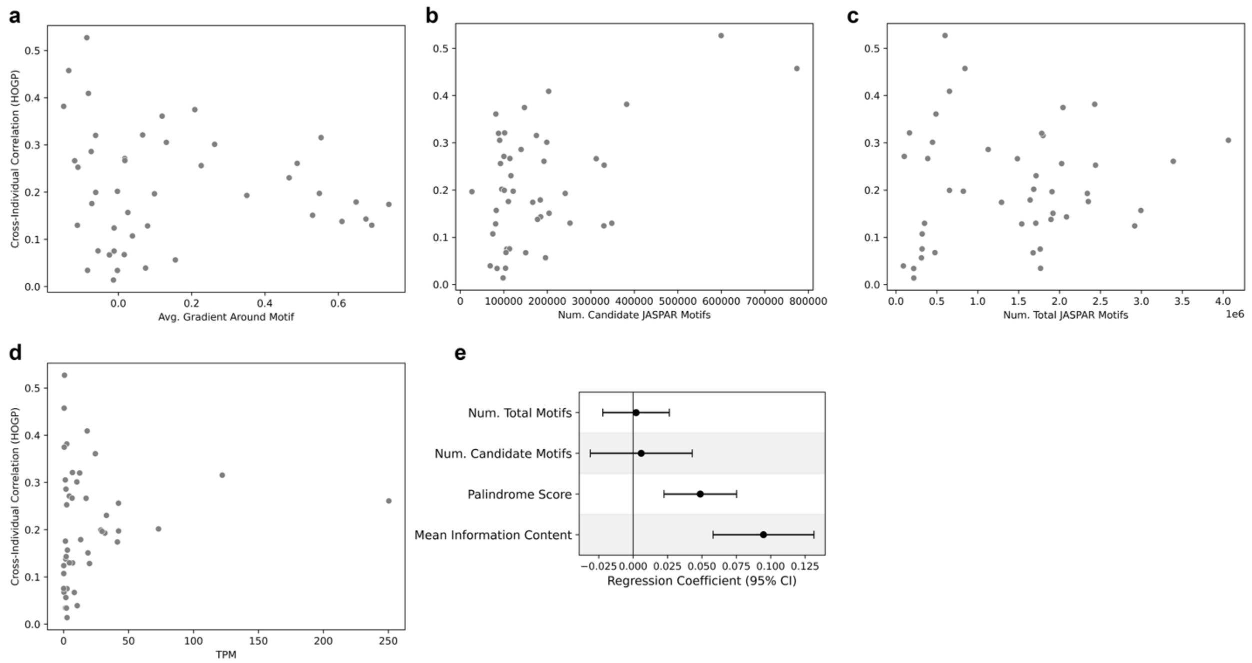


**Supplementary Figure 13: Sequence context variability confounds learning of variant effects independent of motif prevalence, expression, and pre-trained attributions**
(**A**) Cross-individual correlation on HOGP (Y-axis) after fine-tuning Enformer models using aSNVs perturbing one of 45 different motifs, leading to 45 models. On the x-axis is the pre-trained Enformer z-scored gradient attribution (**Methods**) found around all positions of each aSNV-perturbed motif around each of these 1,483 test set genes. The gradient value is averaged over all motif nucleotide positions and each of these genes. Enformer's prior attention paid to each of these motifs is unrelated to its ability to learn new variant effects from within them. The count of candidate (**B,** $p < 10^{-4}$, **Methods**) or all JASPAR-annotated (**C**) motifs found around these 1,483 test set genes is shown on the X-axis instead. We do not observe that fine-tuning on aSNVs that perturb less prevalent motifs lead to more accurate results. In (**D**), we report the expression of the TF linked to each motif on the X-axis, observing no association with HOGP accuracy. In (**E**) we confirm that information content and palindromicity explain variance in HOGP performance independent of motif prevalence via linear regression.

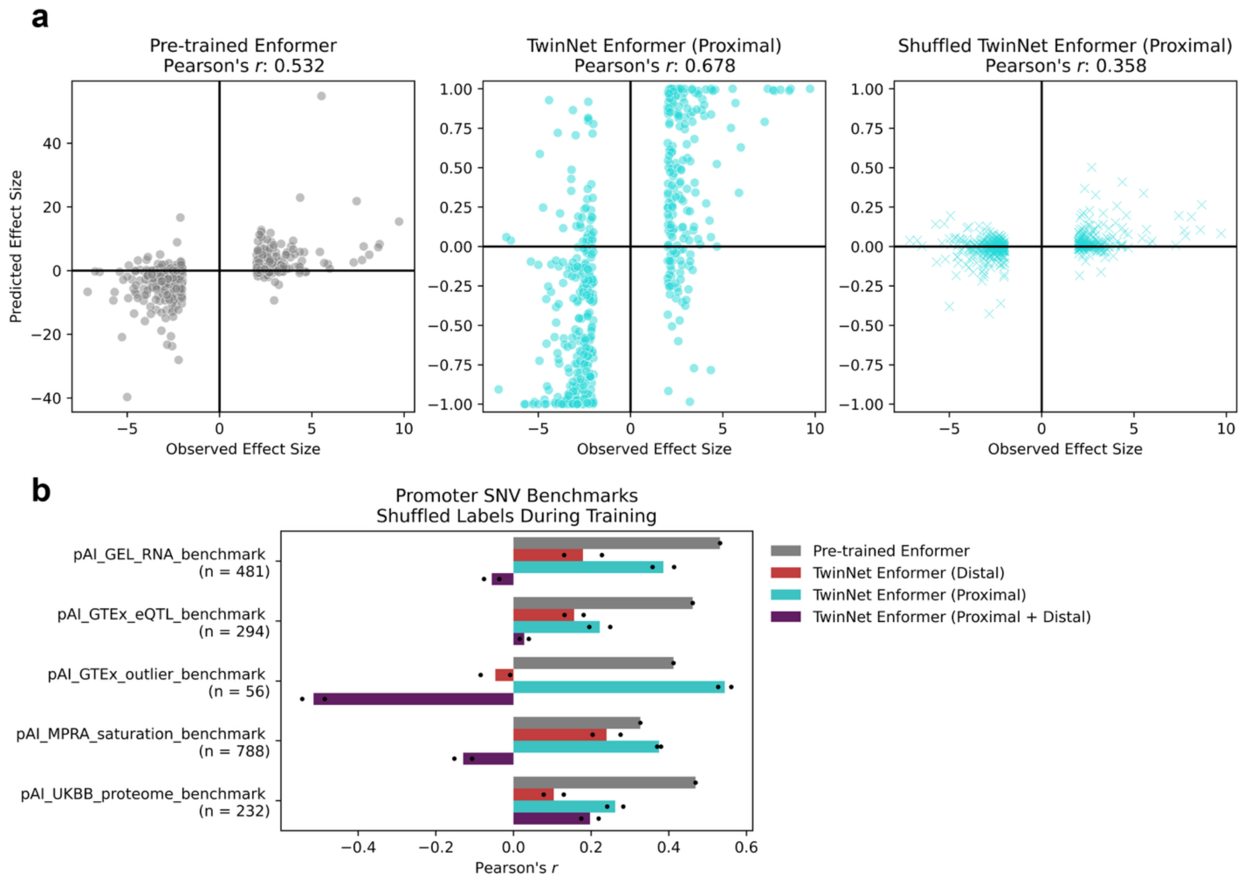


**Supplementary Figure 14: Fine-tuning TwinNet Enformer on rare expression outlier SNVs improves performance on promoter SNV benchmarks**

**A)** Predicted vs observed expression effects for SNVs in the pAI_GEL_RNA_benchmark[15], from Enformer (left), TwinNet Enformer fine-tuned on promoter outliers (middle), or TwinNet Enformer fine-tuned on promoter outliers with the SNV labels shuffled relative to the expression effects (right). **B)** Same as **Fig. 6b**, except each TwinNet Enformer was fine-tuned with SNV labels shuffled relative to expression effects.

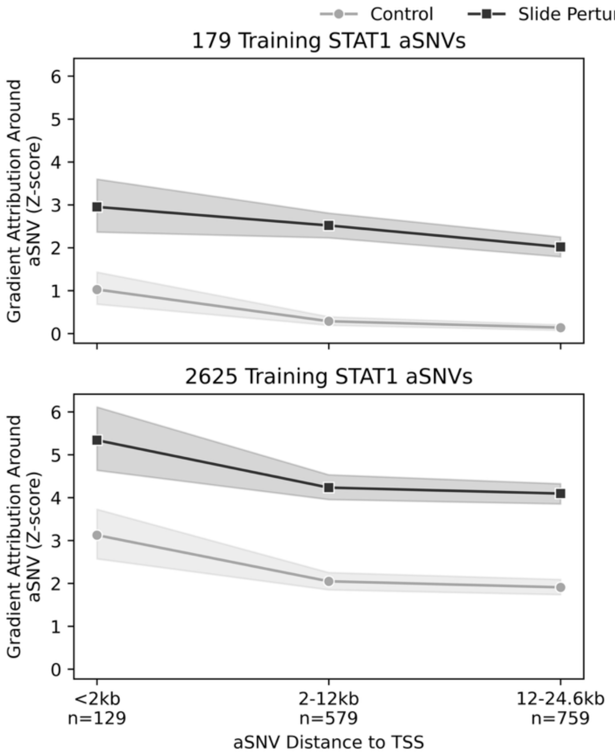


**Supplementary Figure 15: STAT1 gradient attributions after implementing a motif repositioning data augmentation strategy**

(Left**)** Gradients of the gene expression prediction with respect to the input sequence for fine-tuned Enformer models trained with 179 or 2,625 STAT1 aSNVs with (Slide Perturbed Motif Position During Training) or without (Control) the data augmentation strategy from **Fig. 6c**. Gradient values are z-scored such that they represent standard deviations from the mean absolute gradient value per gene (**Methods**). For each of the 1,467 test set genes from **Fig. 6c**, z-scores are averaged over a 21bp window centered on the aSNV-containing STAT1 motif, then grouped by their distance from the gene's TSS. The motif repositioning data augmentation strategy increases gradient mass at all distances. (Right) Gradient values are now normalized relative to the promoter-proximal distance bin (<2kb). Data augmentation leads to less of a dropoff in gradient attribution at greater distances.
